# Learning from imagined experiences via an endogenous prediction error

**DOI:** 10.1101/2024.06.24.600192

**Authors:** Aroma Dabas, Rasmus Bruckner, Heidrun Schultz, Roland G. Benoit

## Abstract

Experiences shape preferences. This is particularly the case when they deviate from our expectations and thus elicit prediction errors. Here we show that prediction errors do not only occur in response to actual events – they also arise endogenously in response to merely imagined events. Specifically, we show that people acquire a preference for acquaintances as they imagine interacting with them in unexpectedly pleasant situations. This learning can best be accounted for by a computational model that calculates prediction errors based on these rewarding experiences. Using functional MRI, we show that the prediction error is mediated via striatal activity. This activity, in turn, seems to update preferences about the individuals by updating their cortical representations. Our findings demonstrate that imaginings can violate our own expectations and thus drive endogenous learning by coopting a neural system that implements reinforcement learning. They reveal fundamental principles how we acquire knowledge devoid of actual experiences.

## Introduction

A hallmark of the human mind is its ability to adapt to an ever-changing environment (1). Our preferences are largely not hard-wired but continuously shaped by experiences. These update the values we assign to objects as well as to the places and people that we encounter in our environment (2). Much of such learning is driven by a mismatch between our expectations and actual experiences (3). For example, when we have an unexpectedly rewarding experience with a particular person, this enhances our expectation that a repeated encounter will also be positive.

Models of reinforcement learning (RL) have formalized this mismatch as a prediction error (PE) (3, 4). This PE serves as a teaching signal that elicits the updating of our values and preferences. We here hypothesize that RL does not only occur in response to external events that we experience (5, 6) or witness (7, 8). Instead, we suggest that it also arises endogenously as a consequence of internal events that we have merely imagined.

The capacity to imagine hypothetical events is often referred to as episodic simulation. It shares several features with episodic memory: The two capacities exhibit parallel developmental trajectories (9, 10), are similarly affected by lesions to the medial temporal lobes (11, 12), and are also more broadly supported by the same core network of brain regions (13). These commonalities have been taken to suggest that episodic simulation is grounded in our memory systems (14, 15). These provide the building blocks for our simulations as well as the constructive processes to recombine these building blocks into novel events.

Given the functional overlap between episodic simulation and memory, we hypothesize that we can also learn from simulated experiences much like we learn from actual experiences. Indeed, motor imagery can enhance motor performance (16) and aversive mental imagery elicits de novo fear conditioning (17). Moreover, when we simply imagine chance encounters with beloved people at random locations, this changes how much we like the location of the imaginary meetings (18, 19).

Such simulation-based learning does not entail any actual feedback from the environment – yet we hypothesize that it induces learning much in the same way as experience-based learning. Episodic simulations can lead to the consideration of alternative outcomes (20, 21) and forge new insights (22, 23). They can thus yield mismatches with our prior beliefs. On an algorithmic level, we hypothesize that such internally generated mismatches cause PE that induces learning in a similar fashion as externally derived mismatches do.

On a neural level, we also hypothesize that this simulation-based RL is based on a mechanism that is shared with experience-based RL. Specifically, the endogenous PE should then also be mediated via dopaminergic activity in the ventral striatum, as demonstrated in neurophysiological recordings in rodents (24, 25) and monkeys (26, 27) and consistent with regional activation observed with human fMRI (28, 29).

We further suggest that this striatal activity enables learning about an environmental stimulus by inducing plasticity in its cortical representation (30, 31). For example, the dorsomedial prefrontal cortex (dmPFC) encodes representations of familiar people (32–34). We would thus expect the ventral striatum to interact with this region, when we experience an unexpected reward while imagining someone that we know.

To test these complementary hypotheses, we developed a procedure that is akin to a stable two-armed bandit task(Fig. 1). In preparation for this task, participants first provided a list of people with whom they are personally familiar. They then rated how much they liked each person. Four of the most neutrally liked people were allocated to a high-reward (HR) and a low-reward (LR) condition.

**Fig. 1.**
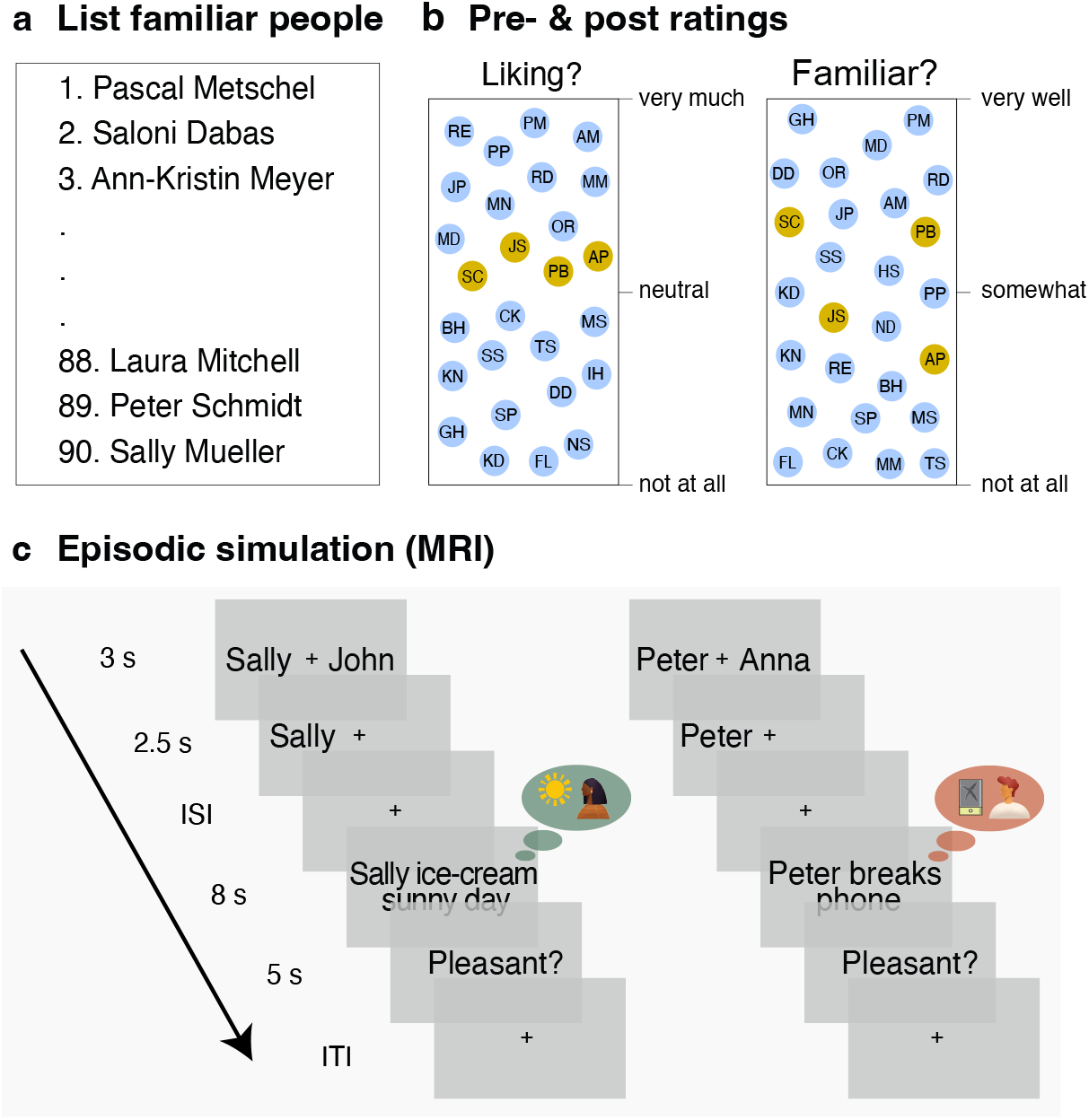
Experimental procedure. **a**, Participants provided names of personally familiar people, and **b**, rated them according to their liking and familiarity - both before and after the episodic simulation task. Based on the initial rating, we selected the six most neutrally liked people, two each to create a high reward (HR), a low reward (LR), and a baseline condition. **c**, On each trial in the MRI scanner, participants made a choice between an HR and an LR person. They sought to select the person that is likely to yield a more pleasant imagined episode. After making their decision, they imagined interacting with the person in either a pleasant (probability ratio: HR/LR = .8/.3) or neutral-to-unpleasant scenario. Participants then rated the pleasantness of the simulated interaction, which served as a proxy for the reward value of the mental experience.

During the fMRI session, participants made repeated choices between two people, one from each reward condition (e.g., Sally vs. Harry). They then imagined a vivid interaction with the chosen person (e.g., Sally) in a specified scenario. These scenarios were either pleasant (e.g., “Sally wishes you well on your birthday”) or neutral to unpleasant (e.g., “Sally returns your bike broken”). The people in the HR condition were imagined in pleasant scenarios with a higher probability than those in the LR condition (80% vs. 30% of the trials). After each trial, participants indicated the pleasantness of the imagined interaction. We take this outcome measure as a proxy for the experienced reward and use it to model the PE on a given trial.

This procedure allowed us to test key features of our hypotheses. First, on a behavioral level, participants seek to maximize the reward that they receive across the experimental session. We thus expected them to acquire a preference for people in the HR, over people in the LR, condition. If this preference shift truly reflects a value update of the people, we also expected a change in how much they like these people as measured on an external rating task. Second, on an algorithmic level, we expected that the preference shift could be accounted for by the Rescorla-Wagner (RW) model (4) of RL.

Third, on a neural level, we expected that this endogenous RL would also be implemented in a similar fashion as experience-based RL. Specifically, in a model-based analysis of our fMRI data, we examined whether the endogenous PE is mediated by neural activity in the ventral striatum. Moreover, using representational similarity and psychophysiological interaction analyses, we tested our account that such a striatal PE updates value by interacting with cortical areas that are involved in representing the information that is being updated. In the case of information about familiar people, such an area should be the dmPFC (32–34).

## Results

### Episodic simulations shift preferences

We first assessed whether participants acquired a preference for selecting people with whom they had mentally experienced more pleasant episodes. To test this prediction, we calculated, for each participant, the overall probability of choosing the HR people across the experimental session. These probabilities were larger than chance, corroborating that episodic simulations indeed induced such a preference (*W* = 1225, *p* < 0.001, *r* = 0.87; Shapiro Wilk: *W* = 0.89, *p* < 0.001; Fig. 2a,b).

**Fig. 2.**
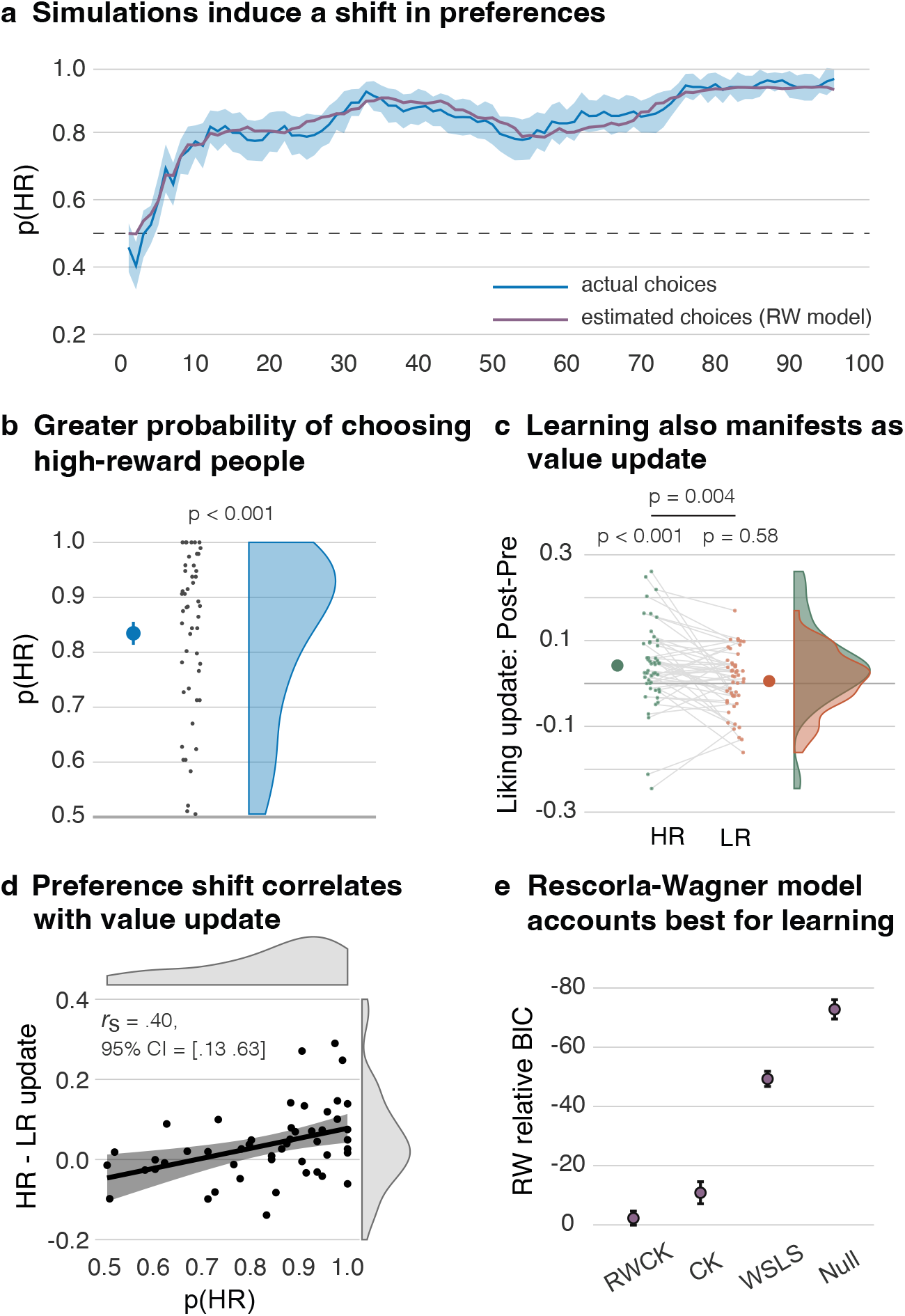
Episodic simulation induced learning. **a**, Over the course of the experiment, participants acquired a preference for selecting people from the high-reward (HR) condition (blue line). Shaded blue area indicates the standard error of mean. The trial-wise estimated choices from the Rescorla-Wagner (RW) model showed a good fit with the empirical data (purple line). **b**, Overall, participants showed a high probability for selecting a HR (versus a low-reward; LR) person. **c**, On the external rating task, episodic simulations also led to an increase in liking for persons in the HR condition that was greater than the absent change in liking for the LR condition. **d**, The acquired preference for HR people in the simulation task (p(HR)) correlated with the increase in liking for HR (vs. LR) people across participants, suggesting that both effects reflect a value update induced by the episodic simulations. Black line shows the linear regression and the shaded area the 95%-confidence intervals. **e**, Relative to the RW model, all other models had a poorer fit to the empirical data. Model fit was estimated using BIC values and the difference was computed as Δ*BIC* = *BIC*_*RW*_ − *BIC*_*model*_. Larger dots denote the means; the error bars the standard error of mean.

### The simulation-induced value update generalizes to an external measure

This preference shift is thought to reflect a positive update of how much the HR people are being valued. If this is the case, we expected the value update to also manifest on an external rating measure. Both before and after the simulation task, participants indicated how much they liked the HR and LR people as well as two further people from a baseline condition that they had not encountered during the simulation task. We could thus examine changes in liking induced by the simulations while controlling for any generic differences across the two measurements. On the initial test, people allocated to the three conditions neither differed in terms of how much they were liked (*F*(2, 46) = 0.03, *p* = 0.97, *n*^2^ < 0.01) nor how familiar they were to the participants (*F*(2, 46) = 0.21, *p* = 0.81, *n*^2^ < 0.01).

To examine the change in liking, we subtracted the liking ratings of the initial test from the one following the simulation task. We then corrected the change scores for the HR and LR conditions by subtracting the change scores for the baseline condition. This measure did not yield a significant change in liking for the LR people (*t*_48_ = 0.54, *p* = 0.58, *d* = 0.08). By contrast, and consistent with our hypothesis, it revealed an increase in liking for the HR people (*W* = 958, *p* < 0.001, *r* = 0.49; Shapiro Wilk: *W* = 0.94, *p* = 0.02). This increase was more pronounced than the absent effect for the LR people (*W* = 875, *p* = 0.004, *r* = 0.37; Shapiro Wilk: *W* = 0.95, *p* = 0.03; Fig. 2c).

If the preference shift and the increase in liking are both manifestations of a simulation-induced value update, we further reasoned that these effects may be correlated with each other across participants. As a measure of acquired preference, we used the probability of choosing the HR people across the experimental session. As a measure of the increase in liking of the HR people, we corrected their liking update (i.e., the post – pre ratings) by subtracting the analogous update for the LR people. By this, both measures examine simulation-induced changes for the HR versus LR condition.

As predicted, these measures were indeed positively associated with each other, as assessed by a robust skipped Spearman’s correlation (*r*_*s*_ = 0.40, 95% CI = [0.13 0.63]; Fig. 2d). (Note that we also obtained a significant effect when performing this analysis with the uncorrected liking update for the HR people; skipped Spearman’s correlation: *r*_*s*_ = 0.38, 95% CI = [0.11 0.61]).

Overall, these results indicate that simulated experiences update our values much like real experiences do. In the next section, we examine the computational and neural processes that guide such learning.

### Reinforcement learning accounts for simulation-based learning

As detailed in the previous sections, episodic simulations induced a shift in preference for people that had been simulated more frequently in rewarding situations. We had hypothesized that such learning is based on an endogenous PE that drives reinforcement learning. To examine this hypothesis, we tested whether the participants’ trial-to-trial adaptations of their preferences can be captured by the RW model.

On each trial, the RW model updates the value of the chosen person as a function of the PE, scaled by the subject-specific learning rate *α* (a free parameter of the model). The PE, in turn, is calculated as the difference between the experienced reward, in this case the pleasantness of the imagined episode, and the current value of the person.

The values are then converted into the probability of selecting a person on a given trial, using a softmax decision rule that contains the freely estimated inverse temperature parameter, *β*. We first determined the individual best fitting *α* and *β* parameters, and then estimated the RW model’s trial-wise choice probabilities and choice values. As evident in Fig. 2a, the model’s estimated choice probability follows closely the trajectory of the participants’ trial-wise choices.

Notably, across participants, a stronger overall probability of the RW model to select the HR people correlated with a greater increase in liking for the HR versus the LR people (see Supplement). Much like the participants’ actual choices, the choice probabilities of the model were thus associated with the external measure of the simulation-induced value update.

We further examined whether the RW model is better at accounting for the choices than a number of competing models of various complexity. Our alternative models explained choices as driven by (a) probabilistically sticking with the rewarded people while avoiding ‘punished’ people (noisy win-stay-lose-shift; WSLS (35)), (b) repeating previous choices (choice kernel; CK (36)), (c) a combination of choice kernel and choice value (RW-CK (36)), and (d) random selection (Null) (for details, see Supplement).

For each model, we used the best-fitting parameter values to compute the Bayesian Information Criterion (BIC) as a measure of the models’ goodness of fit. The simple RW model was indeed the best in accounting for the data (Fig. 2e). We further corroborated this model’s superiority using Bayesian model selection: Model frequencies and exceedance probabilities indicated that the RW model fits the data best of all the included models (Table 1).

**Table 1:**
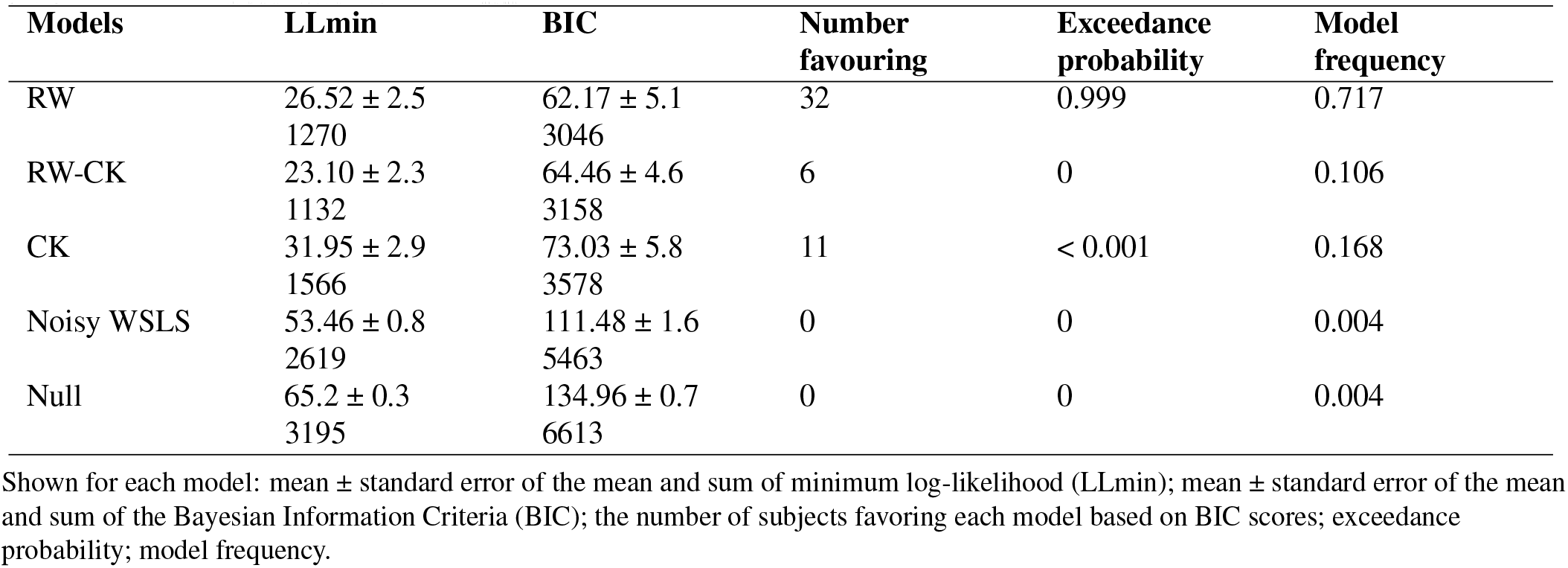
Model comparison.

As hypothesized, simulation-based learning could best be accounted for by a model of reinforcement learning that is driven by an endogenous PE. In the following, we examine whether it is also mediated by the hypothesized brain system.

### The endogenous PE is mediated by the ventral striatum

We first tested the hypothesis that the endogenous PE is mediated by activity in the ventral striatum. Specifically, we conducted a parametric modulation analysis to examine whether trial-by-trial variations in striatal activity can be accounted for by the model-derived time series of the PE. This was indeed the case in our *a priori* region of interest (ROI), an anatomical mask of the ventral striatum (mask from Oxford-GSK-Imanova Structural (37)) (*t*_48_ = 5.66, *p* < 0.001, *d* = 0.81; Fig. 3a).

**Fig. 3.**
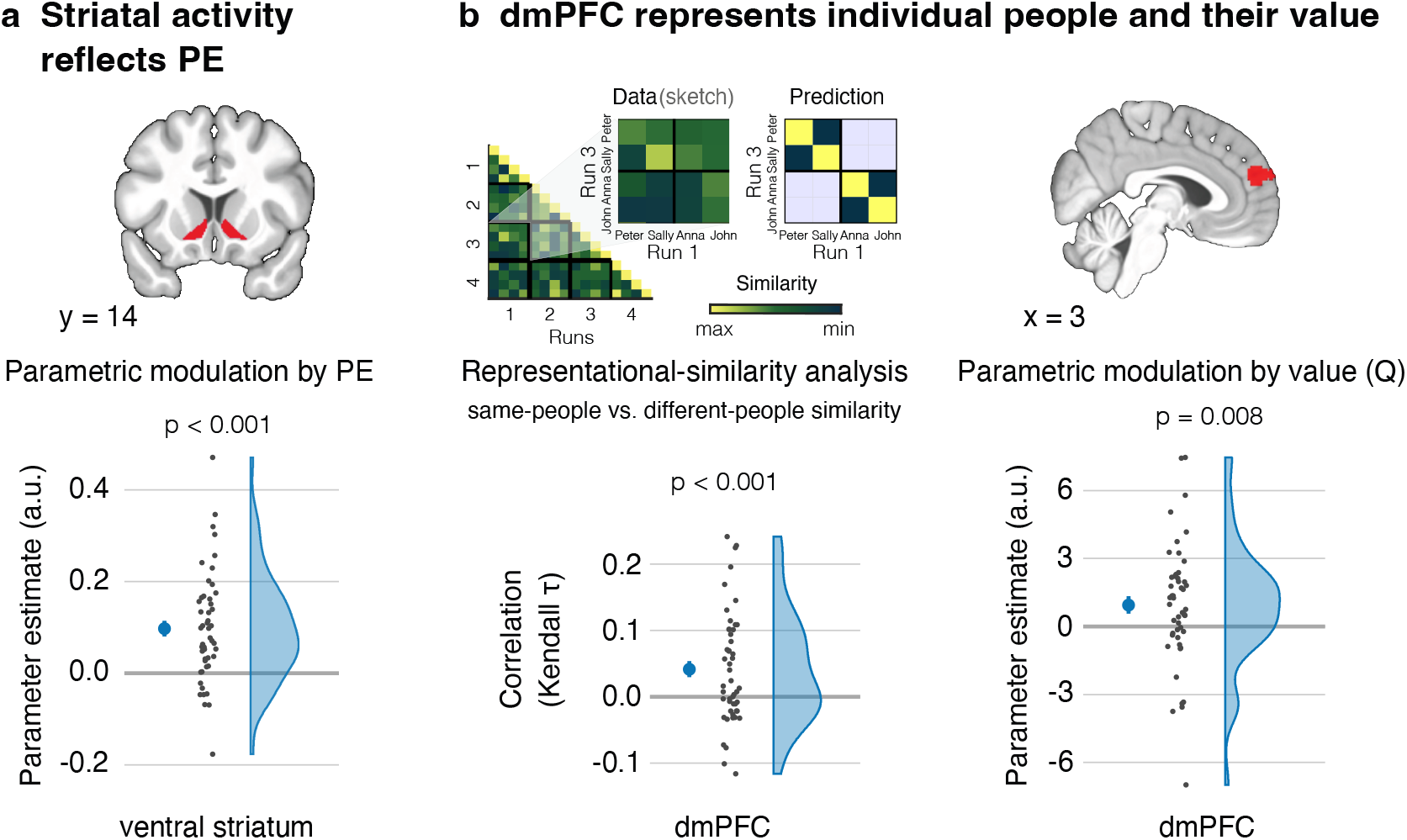
The neural basis of endogenous reinforcement learning. **a**, Activation in the ventral-striatal region-of-interest (ROI) was parametrically modulated by the trial-wise prediction error (PE) as estimated by the Rescorla-Wagner (RW) model. **b**, A representational-similarity analysis indicated that the dorsomedial prefrontal cortex (dmPFC) encodes representation of individual people (left part of panel). This was reflected as more similar activity patterns during the simulation of the same people than of different people across functional runs. Activity in the dmPFC was also parametrically modulated by the continuously updated value of the people (Q) as estimated by the RW model (right part of panel). Activity in the dmPFC thus caries information about the individual people and about their value. Larger dots denote the mean and the errors bars the standard error of mean.

We further corroborated these findings in a complementary whole-brain analysis. As predicted, this analysis yielded a cluster in the bilateral ventral striatum, specifically the nucleus accumbens, that extended into the anterior cingulate cortex (Supplement Table S1 and S2). We also obtained significant clusters in the bilateral anterior hippocampi and parahippocampal cortices, in the paracingulate cortex including the retrosplenial cortex, and the ventromedial prefrontal cortex (Supplement Table S1).

### Striatal-dorsomedial prefrontal interactions support endogenous reinforcement learning

We next sought to test the hypothesis that the ventral striatum updates value by interacting with a cortical region that is involved in representing the information that is being updated. To examine this hypothesis, we first assessed whether the dmPFC encodes representations of individual people and of their value. We focused on this region, given its consistent involvement in thinking about other people (32, 33).

#### Dorsomedial prefrontal cortex codes for representations of people

If the dmPFC encodes representations of individual people, we reasoned that the same representation should get reinstated whenever participants imagine an episode that features the same individual person. We examined this hypothesis by testing the replicability of activity patterns in the dmPFC using representational similarity analysis (38, 39).

Specifically, we employed a region-of-interest that was defined by the *Neurosynth* meta-analysis for the term *people*. In this ROI, we examined whether the activity pattern during a specific episode is more similar to the activity patterns of other episodes that feature the same person (same-person similarity) than a different person (different-person similarity). To avoid any influence of the experimental condition on the results, we calculated the different-person similarity separately for the HR and LR conditions. Consistent with our hypothesis, this effect was significant in the dmPFC (*t*_48_ = 3.47, *p* < 0.001, *d* = 0.50; Fig. 3b, left panel).

We obtained consistent result in an exploratory whole-brain searchlight analysis (see Supplement Table S5 and S6).

#### Dorsomedial prefrontal cortex codes for the persons’ value

The dmPFC thus seems to encode representations of individual people. Does this region also represent their value? To address this question, we took the model-derived value of the chosen person on any trial. This value, the variable Q, is continuously updated by the RW model based on the experienced PE.

We then examined whether regional activation during the choice period is parametrically modulated by Q. Consistent with our prediction, this effect was significant (*t*_48_ = 2.47, *p* = 0.008, *d* = 0.35), indicating that activation in the dmPFC reflects the continuously updated value of the imagined person (Fig. 3b, right panel). Together, the results suggest that the people representations in the dmPFC also entail information about their value.

#### Stronger striatal-dmPFC coupling accompanies stronger value updates

We next examined whether the ventral striatum may elicit the value update by interacting with the dmPFC. Specifically, we conducted a psychophysiological interaction analysis that was seeded in the ventral striatum. The activation time-series was convolved with the trial-specific PE. The ensuing coupling parameters in the dmPFC were significant (*t*_48_ = 4.49, *p* < 0.001, *d* = 0.64), indicating that, as predicted, a stronger value update coincided with a stronger functional connectivity between these regions (Fig. 4).

**Fig. 4.**
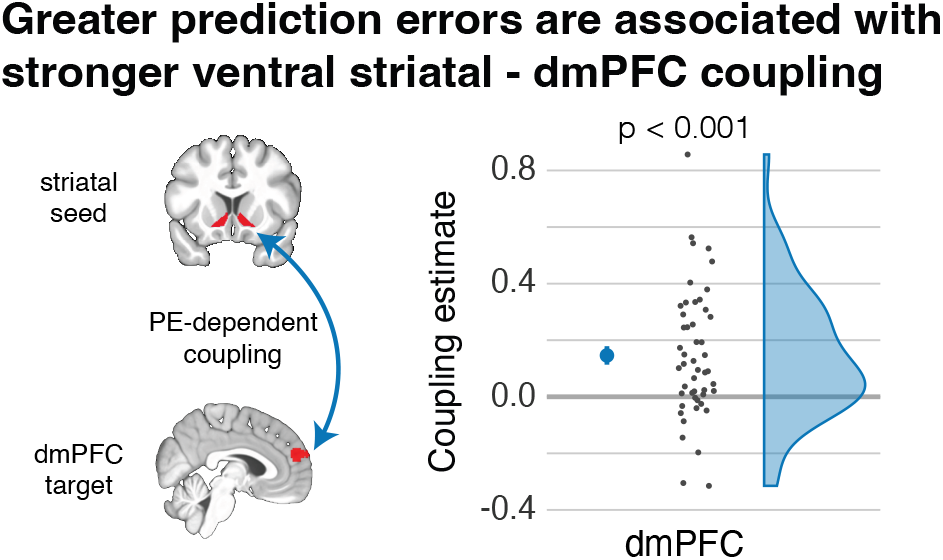
Prediction-error dependent coupling between ventral striatum and dmPFC. A psychophysiological interaction (PPI) analysis revealed that a greater prediction error (PE) is associated with a stronger functional connectivity between the ventral striatum and the dorsomedial prefrontal cortex (dmPFC). It is thus associated with a greater coupling between the region eliciting the value update and the region representing the information that is being updated. The figure displays the PPI effect in the dmPFC. The larger dot denotes the mean and the errors bar the standard error of mean.

## Discussion

To ensure our survival, it is crucial that we continually learn from our past experiences and accurately predict and respond to changes in our environment. Our past experiences play a fundamental role in reinforcing our behavior, guiding us toward rewarding choices, and steering us away from choices that could lead to detrimental outcomes. Notably, our learning is not solely dependent on our direct experiences. We can also learn vicariously by observing the outcomes experienced by others (7, 8, 40).

In this study, we build on this seminal work and demonstrate that merely simulated rewards can induce learning and modify our real-life preferences. Episodic simulations have previously been shown to boost the retention of unpredicted information (41). The current results go beyond these data in demonstrating that the prediction error itself can be a consequence of the simulated experience. Our findings indicate that this simulation-based learning is governed by the same computational and neural mechanisms that underlie direct and vicarious value-based learning.

This is remarkable for at least two reasons. First, simulation-based learning does not entail the actual receipt of a tangible primary (e.g., juice, shock) or secondary (e.g., money) reward. The rewarding outcome in the present study is the pleasantness experienced during the simulation. Yet, it can induce learning in the same fashion. Second, and relatedly, simulation-based learning differs fundamentally from experience-based learning, in that the prediction error does not constitute a mismatch between an internal model of the world and externally-derived information. Instead, the prediction error is based on a mismatch between our internal model *and* internally simulated information. Simulation-based learning is thus driven by an *endogenous* prediction error.

The endogenous prediction error seems to drive learning much like experience-based prediction errors. Specifically, we found that this learning can be described as an effort to minimize these errors as formalized by the Rescorla-Wagner model. This simple model was better at accounting for the data than a number of alternative models of various complexity. Over the years, the Rescorla-Wagner model has been shown to account for diverse phenomena – including Pavlovian and instrumental learning of simple features of the environment up to learning from complex social interactions (42, 43). Here, we generalize those findings by showing that there is no need for an actual reward to trigger learning.

The notion that simulation- and experience-based reinforcement learning are based on a shared mechanism is further supported by our neuroimaging data. These highlight the contribution of the ventral striatum in mediating the endogenous prediction error. This region, and in particular the nucleus accumbens, has consistently been implicated as a key region for signaling PE, irrespective of whether they stem from direct or vicarious learning (28). The present study further highlights the domain-general contribution of this region. In all of these cases, the ventral striatum performs the same computations – only the input that is driving these computations is different.

However, we do not suggest that it is just the striatum that mediates endogenous reinforcement learning. Our exploratory whole-brain analysis identified a network of regions, including the ventral striatum, anterior hippocampus, and ventromedial prefrontal cortex. Activity in all of these regions is typically associated with the magnitude of a prediction error in neuroimaging studies. In particular, the hippocampus has been suggested to play a critical role by detecting novelty and signaling this information to midbrain dopamine neurons via the ventral striatum (44). Striatal (30, 31) and, in particular, dopaminergic activity (24, 45) then induce plasticity in cortical representations and thus affords learning.

In this study, we showed that the dorsomedial prefrontal cortex is involved in representing information about people (32–34) and their continuously updated value. This region showed a stronger coupling with the ventral striatum in case of a stronger prediction error. We thus observed a learning-dependent interaction between the region eliciting the updating of information and the region involved in representing the information that is being updated. These interactions may be mediated by established anatomical connections between these regions, for example through the cortico-basal ganglia-thalamo-cortical circuit (46).

The observation that prediction errors can update our attitudes toward personally familiar people is reminiscent of data demonstrating that a mnemonic prediction error can also update episodic memories (47–50). Specifically, prediction errors can disrupt sustained representations of episodic memories, rendering them malleable and allowing for the integration of new information (47). These data align with the current results in showing that prediction errors can lead to the modification of declarative long-term memory representations.

The demonstration of endogenous reinforcement learning adds to the accumulating evidence that episodic simulation allows us to bootstrap from our experiences to forge new insights. This capacity has both beneficial and unwelcome consequences. On the one hand, episodic simulation constitutes a powerful learning mechanism that doesn’t depend on any new experiences. For example, simulations can help us to come up with solutions to dreaded situations and thus mitigate our apprehensiveness about the future (20, 51). Moreover, simulations allow us to mentally experience how we would feel in prospective episodes (32, 52), and this experience can motivate more farsighted decisions (53, 54).

On the other hand, there is also a downside to a learning mechanism that is decoupled from environmental feedback. In particular, individuals with elevated anxiety or depression show a negative affective bias and have a higher learning rate for negative outcomes (55). There is also some preliminary evidence that people high in neuroticism learn less from imagined positive experiences (18). Without correcting feedback, simulation-based learning may thus contribute to the maintenance of various affective disorders (56, 57). Sometimes it may thus be more beneficial for one’s wellbeing to stop simulating hypothetical events (51, 58, 59).

To conclude, the present study reveals fundamental principles of how we learn from merely imagined experiences. These experiences contrast with our internal expectations and thus induce an endogenous prediction error. Our simulations thus feed into a more general learning mechanism that seeks to reduce uncertainty and that is mediated, among other regions, by the ventral striatum and its interactions with the neocortex. A better understanding of this mechanism may elucidate the origin of a number of maladaptive psychological processes and allow us to also harness its adaptive potential. More broadly, it will be integral for our comprehension of how we create models of our world.

## Methods

### Participants

We recruited 50 participants with no reported history of neurological and psychiatric disorders for the study. They provided written consent as approved by the ethics committee at Leipzig University (ethics number 122/19-ek). One participant fell asleep during the MRI session and was thus excluded from analyses. We accordingly included data from 49 participants (24 female, 25 male; age: *M* = 27.6 y; *SD* = 4.8 y, range = 19 y to 35 y). Prior to data collection, we validated our stimulus material on an independent sample (*n* = 107; 57 female, 50 male; age: *M* = 24.1 y; *SD* = 3.8 y, range = 19 y to 35 y) (see below).

### Materials

This study examined whether episodic simulations induce learning about personally familiar people. Similar to ref. (18, 19), we thus asked participants to compile a list of 90 names of people that they know from their everyday lives (Fig. 1a). They then indicated how much they liked the people. We take liking as a proxy for the degree to which they value a given person. We also assessed how familiar they were with each person. Note that we only acquired these ratings for the respective last 30 people that the participants had provided. This is because people listed earlier tend to be those that are particularly liked, whereas we were interested in selecting more neutral acquaintances that tend to be listed later (18).

Specifically, the participants placed the names of the 30 people on a rectangular space with rating values on the y-axis (Fig. 1b). The y-axis ranged from “not at all liked” to “very much liked” and from “not at all known” to “very well known”. This measurement allowed us to assess the relative liking and familiarity of the people in a fine-grained fashion.

Based on the ratings, we selected six people who were neutrally liked and also sufficiently familiar to the participants. Two of these people were each assigned to the high-reward (HR), low-reward (LR), and baseline conditions, while we aimed to minimize differences in liking and familiarity across conditions.

With an independent sample of participants (*n* = 107), we validated a set of 129 sentences describing scenarios that are either pleasant (67) or neutral-to-unpleasant (62) and that can be imagined in detail (see Supplement for details). The sentences served to induce the simulations of specific episodes in the MRI scanner.

### Experimental procedure

In the MRI scanner, participants performed a simulation task that is akin to a two-armed bandit task (Fig. 1c). Each trial started with a choice phase. Here, they were presented with the names of an HR and an LR person on either side of a fixation cross for a maximum of 3 s. Participants aimed to select the person with whom they would more likely imagine a pleasant experience in the next step.

Once participants had made their choice, only the selected person remained onscreen next to the fixation cross for 2.5 s. This was followed by only the fixation cross for 2.5 s plus any time remaining from the choice phase. Afterward, participants saw the chosen name together with a sentence describing a naturalistic scenario onscreen for 8 s. During this period, they imagined interacting with the person in the presented scenario.

The people in the HR condition were imagined in pleasant scenarios with a higher probability than those in the LR condition (80% vs. 30% of the trials). In the remaining trials, participants imagined interacting with the selected person in one of the neutral-to-unpleasant scenarios.

At the end of each trial, participants indicated the pleasantness of the imagined episode on a continuous slider scale, ranging from “very unpleasant” to “very pleasant” with a “neutral” mid-point, within a maximum of 5 s. We take this outcome measure as a proxy for the experienced reward and use it to model the PE on a given trial. The trial then concluded with a fixation cross during the ITI that was presented for a jittered period ranging from 2–5 s (*M* = 3.3 s, *SD* = 0.83 s) plus the remainder of the time from the preceding rating phase. Overall, the task consisted of 96 trials that were split into four blocks. For another project, the participants also performed a standard bandit task (not reported here).

Outside the scanner, participants once again rated the 30 people on the liking and familiarity scales (post-task ratings). They then completed tasks designed to assess their memory for the simulation task and four questionnaires (a short version of the Big Five Inventory, Beck Depression Inventory II, Mind Wandering Questionnaire, and Vividness of Visual Imagery Questionnaire). The results from the memory tasks and the questionnaires are not reported in this manuscript. To conclude the session, participants were debriefed about the purpose of the study and monetarily reimbursed for their time.

### Computational models

We fitted five models to participants’ trial-wise choices during the simulation task. The model space included (a) a standard RL model that captures learning as driven by PEs, i.e., the Rescorla-Wagner (RW) model; (b) a model that captures the propensity of merely repeating previous choices (Choice Kernel model; CK); (c) a combination of the RW and CK models (RW-CK model); (d) a model that captures the tendency to repeat rewarded actions while shifting away from unrewarded actions (Win-Stay-Lose-Shift; WSLS); and (e) a model that captures no learning (Null model). Below, we provide details of the RW model. Details of the other four models are described in the Supplement.

The RW model updates the expected value (*Q*) of the selected choice (*k*) based on the reward (*r*) received on a given trial (*t*). In our simulation task, the familiar person chosen on a given trial *t* constitutes the choice *k*. We binarized the rated pleasantness, and used it as a proxy for the reward *r*. The learning rule was applied as follows:

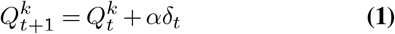

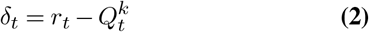

where *α* is the learning rate ranging between 0 and 1 that determines the extent to which the PE at the given trial, *δ*_*t*_, drives the update of the expected value 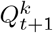. We initialized the choice value 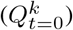 at 0.5, given that the imagined people were considered to be approximately neutral before the start of the task.

We used the softmax choice rule to determine the likelihood that a person is selected on the subsequent trial. The rule uses the computed value of the person from the previous trial as follows:

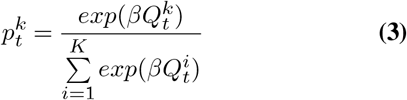

where *β* is the inverse temperature parameter controlling for randomness in choice. It ranges from 0 (completely random) to ∞ (deterministically choosing the most valued person). Overall, the RW model thus has the two free parameters *α* and *β*.

### Model comparison

We sought to assess whether the RW model accounts best for the empirical data by comparing the fits of the different models. To this end, we first fitted each model to participants’ data, by optimizing the model parameters. This was done by minimizing the negative log-likelihood using Matlab’s *fmincon* function.

Using the negative log-likelihoods (*NegLL*), we calculated the Bayesian Information Criteria (*BIC*) for each model and each participant as follows:

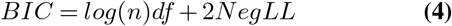

where *df* refers to the number of free parameters and *n* refers to the number of data points. The *BIC* penalizes the maximum likelihood by the number of free parameters. We computed the inter-individual difference in the BIC of all models in relation to the BIC of our hypothesized model, i.e., the RW model (Δ*BIC* = *BIC*_*RW*_ − *BIC*_*model*_) (Fig. 2e).

To assess whether a model fits the data better than all other models in the model set, we analyzed the group-level model fit by estimating the expected frequency and exceedance probability of the model using the VBA toolbox (60) (see Table 1).

Before setting up these analyses on the empirical data, we first ensured that the free parameters of our models could be accurately recovered in simulated data (Supplement and Fig. S2). Moreover, we performed a model recovery analysis based on the simulated data to ensure that we can arbitrate different models (see Supplement and Fig. S3). The analyses were conducted as outlined by ref. (36).

### fMRI data acquisition

Participants were scanned with a 3 Tesla Siemens SKYRA scanner with a 32-channel head coil at the Max Planck Institute for Human Cognitive and Brain Sciences. We acquired structural images with a T1-weighted (T1w) MPRAGE protocol (176 sagittal slices with interleaved acquisition, field of view = 256 mm, 1 mm isotropic voxels, TR = 2300 ms, TE = 5.28 ms, flip angle = 9°, phase encoding: anterior-posterior). For each of the four functional runs, we acquired 307 volumes of blood-oxygen-level-dependent (BOLD) data using a whole brain multiband echo-planar imaging (EPI) sequence (field of view = 204 mm, 2.5 mm isotropic voxels, 60 slices with interleaved acquisition and MF = 3, TR = 2000 ms, TE = 22 ms, flip angle = 80°, phase encoding: anterior-posterior). The first four volumes of each run were discarded to allow for T1w equilibration effects.

### fMRI preprocessing

The MRI data were converted to the Brain Imagining Data Structure (BIDS) (61). The imaging data were then preprocessed using the default preprocessing steps of fMRIprep version 20.2.6 based on Nipype 1.7.0 (62).

The T1w image was corrected for intensity non-uniformity and used as a reference throughout the workflow. The T1w-reference was then skull-tripped and segmented into cerebrospinal fluid (CSF), white matter (WM), and gray matter (GM). It was then normalized to MNI space (MNI152NLin2009cAsym).

The functional imaging data were corrected for slice timing using 0.5 of the slice acquisition range, head motion using the estimated transformation matrices, and the six rotation and translation parameters. They were also corrected for susceptibility distortions using the fieldmap acquired prior to functional MRI imaging. The BOLD reference was co-registered with the T1w reference using boundary-based registration and configured with six degrees of freedom. Several confounding time series were calculated based on the preprocessed BOLD, including the framewise displacement (FD). Additionally, a set of physiological regressors was extracted to allow for component-based noise correction (CompCor). For anatomical CompCor, three probabilistic masks (CSF, WM, and combined CSF+WM) were generated in anatomical space. For further details, please refer to the online documentation (https://fmriprep.org/en/20.2.6/).

The parametric modulation and effective connectivity analyses were performed in MNI space after smoothing with a Gaussian kernel of 6 mm in FWHM. The representational similarity analyses were performed in native space.

### Parametric modulation analysis

The functional MRI data were further analyzed using SPM12 (https://www.fil.ion.ucl.ac.uk/spm/). We performed a parametric modulation analysis to assess activation differences associated with trial-by-trial variations in PE and choice value. Specifically, we estimated a general linear model (GLM) with a boxcar regressor that coded for the 8 s of the simulation period for each trial on which participants had made a choice. This GLM also included regressors coding for the onset of the choice presentations and of the ratings with stick functions. If a subject had missed at least a single trial, we included an additional regressor modeled with a stick function that coded for the onset of each missed trial. Importantly, we included two parametric modulators derived from the RW model: the trial-by-trial PE (*δ*_*t*_; Eqn. 2) modulating the activation during the simulation period and the estimated choice value (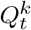 Eqn. 1) during the choice presentation. All of these regressors were convolved with the canonical hemodynamic response function (HRF).

As recommended by ref. (63), we used the population median of the *α* and *β* parameters to estimate the trial-wise PE and Q, given that unregularized random-effects parameter estimates, such as the subject-specific *α* and *β* parameter estimates, tend to be too noisy to obtain reliable neural results.

In our procedure, participants continue to learn the choice value across the whole session. We therefore performed the parametric-modulation analysis on data that were concatenated across the four functional runs. As nuisance regressors, we further included the six head motion parameters, framewise displacement, the first six aCompCor components for each run, and block regressor for the concatenated runs. We adjusted the high-pass filter and temporal non-sphericity calculations to account for the four independent functional runs as implemented in SPM12.

For our region-of-interest (ROI) analyses, we took the mask of the ventral striatum from the FSL Oxford-GSK-Imanova Structural–anatomical Striatal Atlas (Tziortzi et al., 2011) and defined the dorsomedial prefrontal (dmPFC) based on regional cluster obtained from the Neurosynth meta-analysis for the term *people* (association test map). For a given analysis, we then averaged the parameter effect for the respective effect from all voxels within these ROIs.

Additionally, we carried out complementary exploratory whole-brain analyses by entering the respective contrast estimates into a second-level analysis. The results from the whole-brain analyzes are reported in the Supplement.

### Representational similarity analysis

We used representational similarity analysis (RSA) to assess whether the dmPFC region encodes representations of individual people. This analysis used functions from the RSA toolbox (64). We first estimated a GLM that included one boxcar regressor for each imagined person and functional run. These regressors coded for the 8 s of the simulation periods but only for those trials that were congruent to the respective person’s reward category. That is, for HR people, it only included trials with pleasant scenarios; for LR people, it only included trials with neutral-to-unpleasant scenarios.

We then computed the similarity in activity patterns across all voxels of the dmPFC ROI, using Kendall *τ* (64). Specifically, we assessed the similarity of the activity pattern for a given person across runs (*same-person similarity*), and compared this to the similarity of the activity pattern of a given person with the activity pattern for other people across runs (*different-person similarity*) (38). We restricted the different-person similarity to people of the same reward category to ensure that any difference with the same-person similarity is not a result of general condition differences (in particular those related to the value of the imagined event). We complement this anatomical RSA analysis with a spatially unconstrained whole-brain searchlight RSA (see Supplement).

### Psychophysiological connectivity analysis

We computed a psychophysiological interaction (PPI) analysis to test whether the functional connectivity between the ventral striatum (seed region) and dmPFC (target region) varies as a function of the PE. This analysis was based on the GLM from the PE parametric-modulation analysis. The physiological regressor comprised the first eigenvariate of the activity within the ventral striatum, adjusted for effects of interest. The psychological regressor comprised the trial-by-trial PE (*δ*_*t*_; Eqn. 2), estimated using the fixed median *α* and *β* parameters. We computed the PPI regressor by convolving these two regressors, i.e., the activation of the ventral striatum and estimated trial-wise PE. The GLM included these regressors and was estimated with adjusted high-pass filter and temporal non-sphericity calculations to account for the original runs in the concatenated design matrix. To examine changes in coupling with the dmPFC, we extracted the ensuing parameter estimates from our mask for this region.

### Statistical analysis

Statistical tests were conducted with R version 4.2.0 (www.r-project.org). All directed predictions were examined with one-tailed tests. Whenever the Shapiro-Wilk test indicated violations of normality, we used Wilcoxon Signed Rank Test in place of the Student’s t-test. The skipped Spearman’s correlations were computed using the robust correlation toolbox (65) implemented in Matlab (MATLAB version 9.10, R2021a).

### Data and code availability

Pseudonymized behavioral data, list of scenarios, and the custom code for behavioral and fMRI analyses are available on the Open Science Frame-work: https://osf.io/q3v6b/?view_only=625d2433441d41019707903672c71ce5.

The second level t-maps for the parametric-modulation analyses based on PE and Q, searchlight RSA, and psychophysiological interaction analyses, as well as anatomical masks of the dmPFC and ventral striatum ROI, are available at Neurovault: https://identifiers.org/neurovault.collection:17353. Due to identifiable personal markers, we can only share raw MRI images with individual researchers.

## ACKNOWLEDGEMENTS

This work was funded by a Max Planck Research Group awarded to R.G.B. A.D. was also partly supported by the DAAD (Deutscher Akademischer Austauschdienst) and the Max Planck School of Cognition. R.B. was supported by DFG (Deutsche Forschungsgemeinschaft) grant 412917403. We thank Marie-Louise Iredale, Charlotte Gmeiner, Lena Pfund, and Marie-Kristin Mueller for assistance with data collection as well as Maria Woitow for support with developing the event stimuli. We also thank Philipp C. Paulus for sharing his code for the pre- and post-simulation rating task.

## AUTHOR CONTRIBUTIONS

*Conceptualization:* A.D. and R.G.B.; *Methodology:* A.D., and R.G.B.; *Investigation:* A.D.; *Software:* A.D., R.B, and R.G.B.; *Formal Analysis:* A.D., R.B., H.S., and R.G.B.; *Visualization:* A.D.; *Writing – Original Draft:* A.D. and R.G.B.; *Writing – Review & Editing:* A.D., R.B., H.S., and R.G.B.; *Funding Acquisition:* R.G.B.; *Supervision:* R.G.B.

## Bibliography

1. Lucina Q. Uddin. Cognitive and behavioural flexibility: neural mechanisms and clinical considerations. Nat Rev Neurosci, 22(3):167–179, March 2021. ISSN 1471-0048. doi: 10.1038/s41583-021-00428-w. Number: 3 Publisher: Nature Publishing Group.

2. Andreas Olsson, Ewelina Knapska, and Björn Lindström. The neural and computational systems of social learning. Nat Rev Neurosci, 21(4):197–212, April 2020. ISSN 1471-0048. doi: 10.1038/s41583-020-0276-4. Number: 4 Publisher: Nature Publishing Group.

3. Richard S. Sutton and Andrew G. Barto. Reinforcement Learning, second edition: An Introduction. MIT Press, November 2018. ISBN 978-0-262-35270-3. Google-Books-ID: uWV0DwAAQBAJ.

4. RA Rescorla and Allan Wagner. A theory of Pavlovian conditioning: Variations in the effectiveness of reinforcement and nonreinforcement. In Classical Conditioning II: Current Research and Theory, volume Vol. 2. January 1972. Journal Abbreviation: Classical Conditioning II: Current Research and Theory.

5. Timothy E. J. Behrens, Laurence T. Hunt, Mark W. Woolrich, and Matthew F. S. Rushworth. Associative learning of social value. Nature, 456(7219):245–249, November 2008. ISSN 1476-4687. doi: 10.1038/nature07538. Number: 7219 Publisher: Nature Publishing Group.

6. John P. O’Doherty, Peter Dayan, Karl Friston, Hugo Critchley, and Raymond J. Dolan. Temporal difference models and reward-related learning in the human brain. Neuron, 38(2): 329–337, April 2003. ISSN 0896-6273. doi: 10.1016/s0896-6273(03)00169-7.

7. Christopher J. Burke, Philippe N. Tobler, Michelle Baddeley, and Wolfram Schultz. Neural mechanisms of observational learning. Proceedings of the National Academy of Sciences, 107(32):14431–14436, August 2010. doi: 10.1073/pnas.1003111107. Publisher: Proceedings of the National Academy of Sciences.

8. Björn Lindström, Armita Golkar, Simon Jangard, Philippe N. Tobler, and Andreas Olsson. Social threat learning transfers to decision making in humans. Proceedings of the National Academy of Sciences, 116(10):4732–4737, March 2019. doi: 10.1073/pnas.1810180116. Publisher: Proceedings of the National Academy of Sciences.

9. Donna Rose Addis, Alana T. Wong, and Daniel L. Schacter. Age-Related Changes in the Episodic Simulation of Future Events. Psychol Sci, 19(1):33–41, January 2008. ISSN 0956-7976. doi: 10.1111/j.1467-9280.2008.02043.x. Publisher: SAGE Publications Inc.

10. Janie Busby and Thomas Suddendorf. Recalling yesterday and predicting tomorrow. Cognitive Development, 20(3):362–372, July 2005. ISSN 0885-2014. doi: 10.1016/j.cogdev.2005.05.002.

11. Demis Hassabis, Dharshan Kumaran, Seralynne D. Vann, and Eleanor A. Maguire. Patients with hippocampal amnesia cannot imagine new experiences. Proceedings of the National Academy of Sciences, 104(5):1726–1731, January 2007. doi: 10.1073/pnas.0610561104. Publisher: Proceedings of the National Academy of Sciences.

12. Elizabeth Race, Margaret M. Keane, and Mieke Verfaellie. Medial Temporal Lobe Damage Causes Deficits in Episodic Memory and Episodic Future Thinking Not Attributable to Deficits in Narrative Construction. J. Neurosci., 31(28):10262–10269, July 2011. ISSN 0270-6474, 1529-2401. doi: 10.1523/JNEUROSCI.1145-11.2011. Publisher: Society for Neuroscience Section: Articles.

13. Roland G. Benoit and Daniel L. Schacter. Specifying the core network supporting episodic simulation and episodic memory by activation likelihood estimation. Neuropsychologia, 75: 450–457, August 2015. ISSN 0028-3932. doi: 10.1016/j.neuropsychologia.2015.06.034.

14. Daniel L Schacter and Donna Rose Addis. The cognitive neuroscience of constructive memory: remembering the past and imagining the future. Philos Trans R Soc Lond B Biol Sci, 362(1481):773–786, May 2007. ISSN 0962-8436. doi: 10.1098/rstb.2007.2087.

15. Thomas Suddendorf and Michael C. Corballis. The evolution of foresight: What is mental time travel, and is it unique to humans? Behavioral and Brain Sciences, 30(3):299–313, June 2007. ISSN 1469-1825, 0140-525X. doi: 10.1017/S0140525X07001975. Publisher: Cambridge University Press.

16. Rodolphe Gentili, Cheol E. Han, Nicolas Schweighofer, and Charalambos Papaxanthis. Motor Learning Without Doing: Trial-by-Trial Improvement in Motor Performance During Mental Training. Journal of Neurophysiology, 104(2):774–783, August 2010. ISSN 0022-3077. doi: 10.1152/jn.00257.2010. Publisher: American Physiological Society.

17. Erik M. Mueller, Matthias F. J. Sperl, and Christian Panitz. Aversive Imagery Causes De Novo Fear Conditioning. Psychol Sci, 30(7):1001–1015, July 2019. ISSN 0956-7976. doi: 10.1177/0956797619842261. Publisher: SAGE Publications Inc.

18. Philipp C. Paulus, Aroma Dabas, Annalena Felber, and Roland G. Benoit. Simulation-based learning influences real-life attitudes. Cognition, 227:105202, October 2022. ISSN 00100277. doi: 10.1016/j.cognition.2022.105202.

19. Roland G. Benoit, Philipp C. Paulus, and Daniel L. Schacter. Forming attitudes via neural activity supporting affective episodic simulations. Nature Communications, 10(1):2215, May 2019. ISSN 2041-1723. doi: 10.1038/s41467-019-09961-w. Number: 1 Publisher: Nature Publishing Group.

20. Helen G. Jing, Kevin P. Madore, and Daniel L. Schacter. Worrying about the future: An episodic specificity induction impacts problem solving, reappraisal, and well-being. Journal of Experimental Psychology: General, 145(4):402–418, April 2016. ISSN 1939-2222, 0096-3445. doi: 10.1037/xge0000142.

21. Daniel L. Schacter, Roland G. Benoit, Felipe De Brigard, and Karl K. Szpunar. Episodic future thinking and episodic counterfactual thinking: Intersections between memory and decisions. Neurobiology of Learning and Memory, 117:14–21, January 2015. ISSN 1074-7427. doi: 10.1016/j.nlm.2013.12.008.

22. Donna Rose Addis, Ling Pan, Regina Musicaro, and Daniel L. Schacter. Divergent thinking and constructing episodic simulations. Memory, 24(1):89–97, January 2016. ISSN 0965-8211. doi: 10.1080/09658211.2014.985591. Publisher: Routledge _eprint: 10.1080/09658211.2014.985591.

23. Mathias Benedek, Roger E. Beaty, Daniel L. Schacter, and Yoed N. Kenett. The role of memory in creative ideation. Nat Rev Psychol, 2(4):246–257, April 2023. ISSN 2731-0574. doi: 10.1038/s44159-023-00158-z. Number: 4 Publisher: Nature Publishing Group.

24. John N.J Reynolds and Jeffery R Wickens. Dopamine-dependent plasticity of corticostriatal synapses. Neural Networks, 15(4-6):507–521, June 2002. ISSN 08936080. doi: 10.1016/S0893-6080(02)00045-X.

25. Yuji K. Takahashi, Angela J. Langdon, Yael Niv, and Geoffrey Schoenbaum. Temporal Specificity of Reward Prediction Errors Signaled by Putative Dopamine Neurons in Rat VTA Depends on Ventral Striatum. Neuron, 91(1):182–193, July 2016. ISSN 08966273. doi: 10.1016/j.neuron.2016.05.015.

26. J. R. Hollerman and W. Schultz. Dopamine neurons report an error in the temporal prediction of reward during learning. Nat Neurosci, 1(4):304–309, August 1998. ISSN 1097-6256. doi: 10.1038/1124.

27. Wolfram Schultz, Léon Tremblay, and Jeffrey R Hollerman. Reward prediction in primate basal ganglia and frontal cortex. Neuropharmacology, 37(4):421–429, April 1998. ISSN 0028-3908. doi: 10.1016/S0028-3908(98)00071-9.

28. Philip R. Corlett, Jessica A. Mollick, and Hedy Kober. Meta-analysis of human prediction error for incentives, perception, cognition, and action. Neuropsychopharmacol., 47(7): 1339–1349, June 2022. ISSN 1740-634X. doi: 10.1038/s41386-021-01264-3. Number: 7 Publisher: Nature Publishing Group.

29. Mathias Pessiglione, Ben Seymour, Guillaume Flandin, Raymond J. Dolan, and Chris D. Frith. Dopamine-dependent prediction errors underpin reward-seeking behaviour in humans. Nature, 442(7106):1042–1045, August 2006. ISSN 1476-4687. doi: 10.1038/nature05051. Number: 7106 Publisher: Nature Publishing Group.

30. Hanneke E. M. den Ouden, Jean Daunizeau, Jonathan Roiser, Karl J. Friston, and Klaas E. Stephan. Striatal Prediction Error Modulates Cortical Coupling. J. Neurosci., 30(9):3210–3219, March 2010. ISSN 0270-6474, 1529-2401. doi: 10.1523/JNEUROSCI.4458-09.2010. Publisher: Society for Neuroscience Section: Articles.

31. Mona M. Garvert, Michael Moutoussis, Zeb Kurth-Nelson, Timothy E. J. Behrens, and Raymond J. Dolan. Learning-Induced Plasticity in Medial Prefrontal Cortex Predicts Preference Malleability. Neuron, 85(2):418–428, January 2015. ISSN 0896-6273. doi: 10.1016/j.neuron.2014.12.033.

32. Roland G. Benoit, Karl K. Szpunar, and Daniel L. Schacter. Ventromedial prefrontal cortex supports affective future simulation by integrating distributed knowledge. Proc Natl Acad Sci USA, 111(46):16550–16555, November 2014. ISSN 0027-8424, 1091-6490. doi: 10.1073/pnas.1419274111.

33. Demis Hassabis, R. Nathan Spreng, Andrei A. Rusu, Clifford A. Robbins, Raymond A. Mar, and Daniel L. Schacter. Imagine All the People: How the Brain Creates and Uses Personality Models to Predict Behavior. Cerebral Cortex, 24(8):1979–1987, August 2014. ISSN 1047-3211. doi: 10.1093/cercor/bht042.

34. Karl K. Szpunar, Peggy L. St. Jacques, Clifford A. Robbins, Gagan S. Wig, and Daniel L. Schacter. Repetition-related reductions in neural activity reveal component processes of mental simulation. Social Cognitive and Affective Neuroscience, 9(5):712–722, May 2014. ISSN 1749-5016. doi: 10.1093/scan/nst035.

35. John W. Cotton and Chester W. Harris. Reliability coefficients as a function of individual differences induced by a learning process assuming identical organisms. Journal of Mathematical Psychology, 10(4):387–420, November 1973. ISSN 0022-2496. doi: 10.1016/0022-2496(73)90024-2.

36. Robert C Wilson and Anne GE Collins. Ten simple rules for the computational modeling of behavioral data. eLife, 8:e49547, November 2019. ISSN 2050-084X. doi: 10.7554/eLife.49547. Publisher: eLife Sciences Publications, Ltd.

37. Andri C. Tziortzi, Graham E. Searle, Sofia Tzimopoulou, Cristian Salinas, John D. Beaver, Mark Jenkinson, Marc Laruelle, Eugenii A. Rabiner, and Roger N. Gunn. Imaging dopamine receptors in humans with [11C]-(+)-PHNO: dissection of D3 signal and anatomy. Neuroimage, 54(1):264–277, January 2011. ISSN 1095-9572. doi: 10.1016/j.neuroimage.2010.06.044.

38. Hamed Nili, Alexander Walther, Arjen Alink, and Nikolaus Kriegeskorte. Inferring exemplar discriminability in brain representations. PLOS ONE, 15(6):e0232551, June 2020. ISSN 1932-6203. doi: 10.1371/journal.pone.0232551. Publisher: Public Library of Science.

39. Philipp C. Paulus, Ian Charest, and Roland G. Benoit. Value shapes the structure of schematic representations in the medial prefrontal cortex. preprint, Neuroscience, August 2020.

40. Shinsuke Suzuki, Norihiro Harasawa, Kenichi Ueno, Justin L. Gardner, Noritaka Ichinohe, Masahiko Haruno, Kang Cheng, and Hiroyuki Nakahara. Learning to Simulate Others’ Decisions. Neuron, 74(6):1125–1137, June 2012. ISSN 0896-6273. doi: 10.1016/j.neuron.2012.04.030.

41. Alyssa H. Sinclair, Shabnam Hakimi, Matthew L. Stanley, R. Alison Adcock, and Gregory R. Samanez-Larkin. Pairing facts with imagined consequences improves pandemic-related risk perception. Proceedings of the National Academy of Sciences, 118(32):e2100970118, August 2021. doi: 10.1073/pnas.2100970118. Publisher: Proceedings of the National Academy of Sciences.

42. Timothy E. J. Behrens, Laurence T. Hunt, and Matthew F. S. Rushworth. The Computation of Social Behavior. Science, 324(5931):1160–1164, May 2009. doi: 10.1126/science.1169694. Publisher: American Association for the Advancement of Science.

43. Lei Zhang, Lukas Lengersdorff, Nace Mikus, Jan Gläscher, and Claus Lamm. Using reinforcement learning models in social neuroscience: frameworks, pitfalls and suggestions of best practices. Social Cognitive and Affective Neuroscience, 15(6):695–707, July 2020. ISSN 1749-5016. doi: 10.1093/scan/nsaa089.

44. John E. Lisman and Anthony A. Grace. The hippocampal-VTA loop: controlling the entry of information into long-term memory. Neuron, 46(5):703–713, June 2005. ISSN 0896-6273. doi: 10.1016/j.neuron.2005.05.002.

45. Shaowen Bao, Vincent T. Chan, and Michael M. Merzenich. Cortical remodelling induced by activity of ventral tegmental dopamine neurons. Nature, 412(6842):79–83, July 2001. ISSN 1476-4687. doi: 10.1038/35083586. Number: 6842 Publisher: Nature Publishing Group.

46. Tiago V. Maia and Michael J. Frank. From reinforcement learning models to psychiatric and neurological disorders. Nat Neurosci, 14(2):154–162, February 2011. ISSN 1546-1726. doi: 10.1038/nn.2723. Number: 2 Publisher: Nature Publishing Group.

47. Alyssa H. Sinclair, Grace M. Manalili, Iva K. Brunec, R. Alison Adcock, and Morgan D. Barense. Prediction errors disrupt hippocampal representations and update episodic memories. Proceedings of the National Academy of Sciences, 118(51):e2117625118, December 2021. doi: 10.1073/pnas.2117625118. Publisher: Proceedings of the National Academy of Sciences.

48. Oded Bein, Camille Gasser, Tarek Amer, Anat Maril, and Lila Davachi. Predictions transform memories: How expected versus unexpected events are integrated or separated in memory. Neuroscience & Biobehavioral Reviews, 153:105368, October 2023. ISSN 01497634. doi: 10.1016/j.neubiorev.2023.105368.

49. Francesco Pupillo and Rasmus Bruckner. Signed and unsigned effects of prediction error on memory: Is it a matter of choice? Neuroscience & Biobehavioral Reviews, 153:105371, October 2023. ISSN 0149-7634. doi: 10.1016/j.neubiorev.2023.105371.

50. Nina Rouhani, Yael Niv, Michael J. Frank, and Lars Schwabe. Multiple routes to enhanced memory for emotionally relevant events. Trends in Cognitive Sciences, 27(9):867–882, September 2023. ISSN 1364-6613, 1879-307X. doi: 10.1016/j.tics.2023.06.006. Publisher: Elsevier.

51. Roland G. Benoit, Daniel J. Davies, and Michael C. Anderson. Reducing future fears by suppressing the brain mechanisms underlying episodic simulation. Proceedings of the National Academy of Sciences, 113(52):E8492–E8501, December 2016. doi: 10.1073/pnas.1606604114. Publisher: Proceedings of the National Academy of Sciences.

52. Chantelle M. Cocquyt and Daniela J. Palombo. Emotion in the mind’s eye: Imagination for adaptive cognition. Annals of the New York Academy of Sciences, 1526(1):59–72, 2023. ISSN 1749-6632. doi: 10.1111/nyas.15011. _eprint: https://onlinelibrary.wiley.com/doi/pdf/10.1111/nyas.15011.

53. Sarah A. Rösch, Davide F. Stramaccia, and Roland G. Benoit. Promoting farsighted decisions via episodic future thinking: A meta-analysis. Journal of Experimental Psychology: General, 151(7):1606, 2022. Publisher: American Psychological Association.

54. Pascal Boyer. Evolutionary economics of mental time travel? Trends in Cognitive Sciences, 12(6):219–224, June 2008. ISSN 1364-6613. doi: 10.1016/j.tics.2008.03.003.

55. Alexandra C. Pike and Oliver J. Robinson. Reinforcement Learning in Patients With Mood and Anxiety Disorders vs Control Individuals: A Systematic Review and Meta-analysis. JAMA Psychiatry, 79(4):313–322, April 2022. ISSN 2168-622X. doi: 10.1001/jamapsychiatry.2022.0051.

56. Adam Bulley, Julie D. Henry, and Thomas Suddendorf. Thinking about threats: Memory and prospection in human threat management. Consciousness and Cognition, 49:53–69, March 2017. ISSN 1053-8100. doi: 10.1016/j.concog.2017.01.005.

57. Fritz Renner, Julie L. Ji, Arnaud Pictet, Emily A. Holmes, and Simon E. Blackwell. Effects of Engaging in Repeated Mental Imagery of Future Positive Events on Behavioural Activation in Individuals with Major Depressive Disorder. Cognit Ther Res, 41(3):369–380, 2017. ISSN 0147-5916. doi: 10.1007/s10608-016-9776-y.

58. Stephanie M. Ashton, Tom Smeets, and Conny W. E. M. Quaedflieg. Controlling intrusive thoughts of future fears under stress. Neurobiology of Stress, 27:100582, November 2023. ISSN 2352-2895. doi: 10.1016/j.ynstr.2023.100582.

59. Zulkayda Mamat and Michael C. Anderson. Improving mental health by training the suppression of unwanted thoughts. Sci Adv, 9(38):eadh5292, September 2023. ISSN 2375-2548. doi: 10.1126/sciadv.adh5292.

60. Jean Daunizeau, Vincent Adam, and Lionel Rigoux. VBA: A Probabilistic Treatment of Non-linear Models for Neurobiological and Behavioural Data. PLOS Computational Biology, 10 (1):e1003441, January 2014. ISSN 1553-7358. doi: 10.1371/journal.pcbi.1003441. Publisher: Public Library of Science.

61. Krzysztof J. Gorgolewski, Tibor Auer, Vince D. Calhoun, R. Cameron Craddock, Samir Das, Eugene P. Duff, Guillaume Flandin, Satrajit S. Ghosh, Tristan Glatard, Yaroslav O. Halchenko, Daniel A. Handwerker, Michael Hanke, David Keator, Xiangrui Li, Zachary Michael, Camille Maumet, B. Nolan Nichols, Thomas E. Nichols, John Pellman, Jean-Baptiste Poline, Ariel Rokem, Gunnar Schaefer, Vanessa Sochat, William Triplett, Jessica A. Turner, Gaël Varoquaux, and Russell A. Poldrack. The brain imaging data structure, a format for organizing and describing outputs of neuroimaging experiments. Sci Data, 3(1): 160044, June 2016. ISSN 2052-4463. doi: 10.1038/sdata.2016.44. Number: 1 Publisher: Nature Publishing Group.

62. Oscar Esteban, Christopher J. Markiewicz, Ross W. Blair, Craig A. Moodie, A. Ilkay Isik, Asier Erramuzpe, James D. Kent, Mathias Goncalves, Elizabeth DuPre, Madeleine Snyder, Hiroyuki Oya, Satrajit S. Ghosh, Jessey Wright, Joke Durnez, Russell A. Poldrack, and Krzysztof J. Gorgolewski. fMRIPrep: a robust preprocessing pipeline for functional MRI. Nat Methods, 16(1):111–116, December 2018. ISSN 1548-7105. doi: 10.1038/s41592-018-0235-4. Number: 1 Publisher: Nature Publishing Group.

63. Samuel J. Gershman, Bijan Pesaran, and Nathaniel D. Daw. Human Reinforcement Learning Subdivides Structured Action Spaces by Learning Effector-Specific Values. J Neurosci, 29(43):13524–13531, October 2009. ISSN 0270-6474. doi: 10.1523/JNEUROSCI.2469-09.2009.

64. Hamed Nili, Cai Wingfield, Alexander Walther, Li Su, William Marslen-Wilson, and Nikolaus Kriegeskorte. A Toolbox for Representational Similarity Analysis. PLOS Computational Biology, 10(4):e1003553, April 2014. ISSN 1553-7358. doi: 10.1371/journal.pcbi.1003553. Publisher: Public Library of Science.

65. Cyril Pernet, Rand Wilcox, and Guillaume Rousselet. Robust Correlation Analyses: False Positive and Power Validation Using a New Open Source Matlab Toolbox. Frontiers in Psychology, 3, 2013. ISSN 1664-1078.

