## Supplement Document for "Learning from imagined experiences via an endogenous prediction error"

#### 1 Scenario selection

We created a corpus of 129 sentences describing 67 pleasant and 62 neutral-to-unpleasant scenarios. We validated this categorization in a separate sample of participants ($n$ = 107; 57 female, 50 male; age: $M$ = 24.1 y; $SD$ = 3.8 y, range = 19 y to 35 y). Similar to our main experiment, we first requested the participants to provide names of people that they were personally familiar with, and then identified four individuals who they felt neutral towards. Participants were then presented with a scenario and a given individual. They were instructed to immediately immerse themselves in the scenario and to imagine this scenario together with the respective person as vividly as possible. After simulating the interaction for 8 s, participants were prompted to assess the pleasantness of the interaction within 5 s using a continuous scale ranging from "very unpleasant" to "very pleasant," with "neutral" positioned at the midpoint.

We then compared pleasantness ratings between the two scenario conditions while controlling for participant variability using a linear mixed-effects model. Our analysis

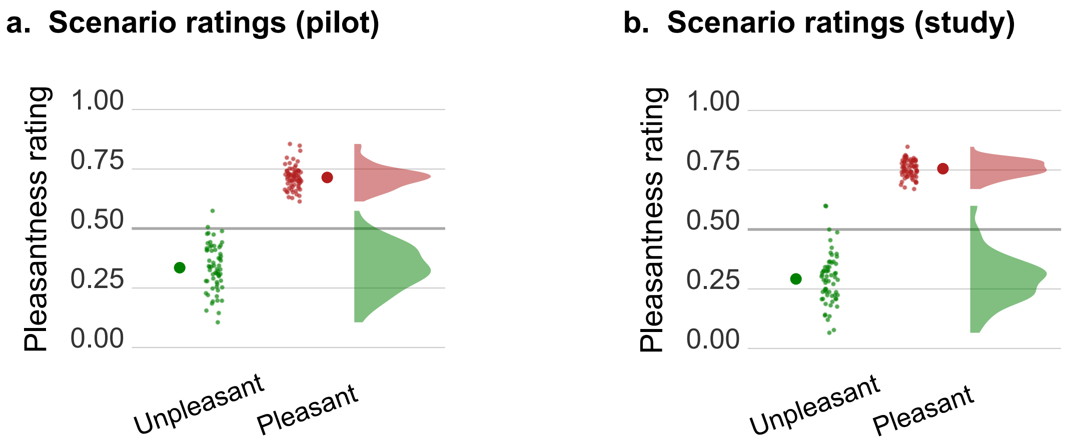


Figure S1: **Scenario ratings from a,** pilot study (n = 107) and **b,** main study (n = 49). Smaller dots denote mean ratings for each scenario. Larger dots denote mean ratings for the conditions.

corroborated that the scenarios categorized as pleasant elicited significantly higher pleasantness ratings compared to the scenarios categorizes as neutral-to-unpleasant (*B* = 0.37, *SE* = 4.5e-03, *z* = 82.48, *p* < 0.001; Fig. S1a). We further confirmed that this was also the case in the main study (*B* = 0.46, *SE* = 5.0e-03, *z* = 91.48, *p* < 0.001; Fig. S1b).

### 2 Computational models

#### 2.1 Model description

Our key model of interest, i.e. the Rescorla-Wagner (RW) model, has been described in the methods section of the main text. Below, we outline the details of the four additional models that capture different decision-making policies.

##### 2.1.1 Choice Kernel (CK) model

The CK model captures behavior that merely repeats previous choices. Specifically, this model tracks the frequency with which all choices were previously selected, in the form of a 'choice kernel', and uses the information to predict choice repetition. The model does not take into account the reward experienced during the simulation. It is implemented as follows:

$$CK_{t+1}^k = CK_t^k + \alpha_c(a_t^k - CK_t^k) \tag{S1}$$

where $CK_t^k$ denotes the choice kernel on trial $t$ and for option $k$. $\alpha_c$ refers to the choice kernel learning rate and $a_t^k$ denotes choice repetition such that $a_t^k$ = 1 if option $k$ is selected on trial $t$, otherwise $a_t^k$ = 0. The value of $\alpha_c$ ranges from 0 to 1, controlling the degree to which the choice repetition (or switch) updates the kernel. We assume that the initial value of the choice kernel for all options is 0. To convert the kernel into choice probabilities, we use the softmax decision rule:

$$p_t^k = \frac{exp(\beta_c CK_t^k)}{\sum\limits_{i=1}^{K} exp(\beta_c CK_t^i)} \tag{S2}$$

where $\beta$ corresponds to the inverse temperature associated with the choice kernel. Overall, this model has the two free parameters $\alpha_c$ and $\beta_c$.

###### 2.1.2 Combined Rescorla-Wagner and Choice Kernel (RW-CK) model

The combined Rescorla-Wagner and choice kernel (RW-CK) model captures choice behavior based on both the value update (which updates according to equation 1 in the main manuscript) and the choice kernel (which updates as per equation S1 in this document). The choice probability is computed by combining the outputs of the RW model and the CK model as follows:

$$p_t^k = \frac{exp(\beta Q_t^k + \beta_c CK_t^k)}{\sum_{i=1}^{K} exp(\beta Q_t^i + \beta_c CK_t^i)} \tag{S3}$$

where $\beta$ and $\beta_c$ denote the inverse temperature parameter associated with choice value $Q_t^i$ and choice kernel $CK_t^i$, respectively. In addition to the inverse temperature parameters, this model has two more free parameters: The learning rates $\alpha$ (Eqn. 1) and $\alpha_c$ (Eqn. S1) of choice value and kernel, respectively.

###### 2.1.3 Win-Stay-Lose-Shift (WSLS) model

This model adapts its behavior according to previous feedback, favoring actions that have been rewarded and avoiding those that haven't. This strategy, known as the win-stay lose-shift rule, is applied with a probability of $1 - \epsilon/2$ , where $\epsilon/2$ captures choice stochasticity. The total probability of staying with option $k$ at time $t$ is thus the combination of two cases: staying after a win with probability $\epsilon/2$ and staying after a loss with probability $\epsilon/2$:

$$p_t^k = \begin{cases} 1 - \epsilon/2 \text{ if } (c_{t-1} = k \text{ and } r_{t-1} = 1) \text{ or } (c_{t-1} \neq k \text{ and } r_{t-1} = 0) \\ \epsilon/2 \text{ if } (c_{t-1} \neq k \text{ and } r_{t-1} = 1) \text{ or } (c_{t-1} = k \text{ and } r_{t-1} = 0) \end{cases} \tag{S4}$$

In the WSLS model, $\epsilon$ is the only free parameter.

###### 2.1.4 Null model

This model captures random choice behavior. The probability of selecting one of the two presented options is fixed at 0.5. This model thus has no free parameter.

#### 2.2 Parameter recovery

Before collecting data, we tested whether our fitting procedure can return meaningful parameter values. We followed the approach outlined by Wilson & Collins (2019). We first simulated data with known parameter values and then estimated the parameters that best describe the simulated data. In an ideal situation, the estimation procedure should yield the parameter value that was used for generating the data.

We ran the fitting procedure 50 times with different random initial conditions to increase the likelihood of finding global minima. Post data collection, we updated the parameter space of the models to match the range recovered from fitting participants' data. The difference between the true and estimated parameter values is plotted in Figure S2.

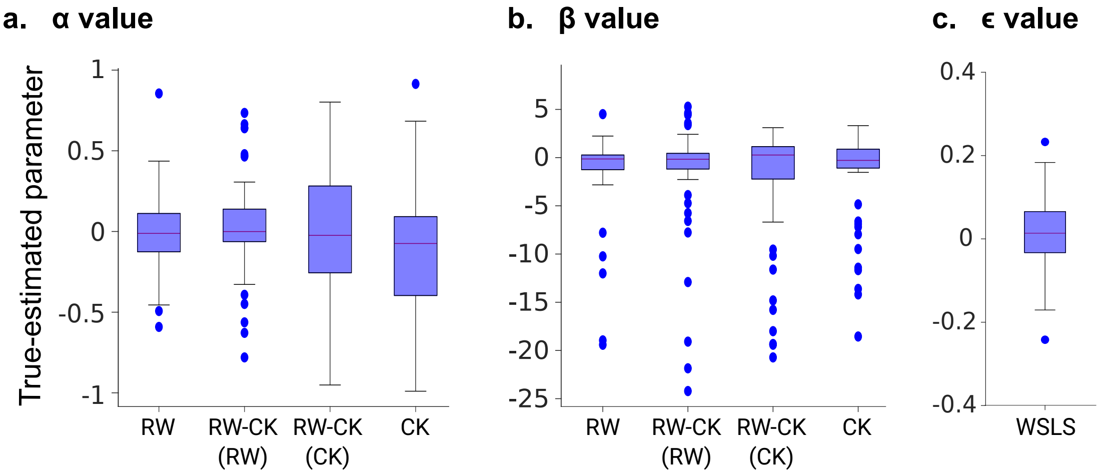


Figure S2: **The difference in the estimated parameter value from its true parameter value.** The median of the boxplots lies close to the 0 value indicating that overall the fitting procedure estimates parameters accurately and without a systematic estimation bias. However, the variability of the estimated parameters is considerably larger for the CK components compared to the RW components. The recovery is based on 50 simulated data sets. The whiskers extend to 1.5 x interquartile range. RW: Rescorla-Wagner model; RW-CK (RW): Rescorla-Wagner part of the combined Rescorla-Wagner Choice-Kernel model; RW-CK (CK): Choice-Kernel part of the RW-CK model; CK: Choice-Kernel; WSLS: Win-Stay-Lose-Shift model.

#### 2.3 Model recovery

We also tested whether our fitting procedure can arbitrate different models. First, we tested whether the model that generates the data is also the model that best fits the data. We illustrate this through the use of confusion matrices (Fig. S3a and S3b). A confusion matrix calculates the probability that the data generated by one model is best fit by a given model, i.e. p(fit model = B | simulated model = A), considering all models within the model space.

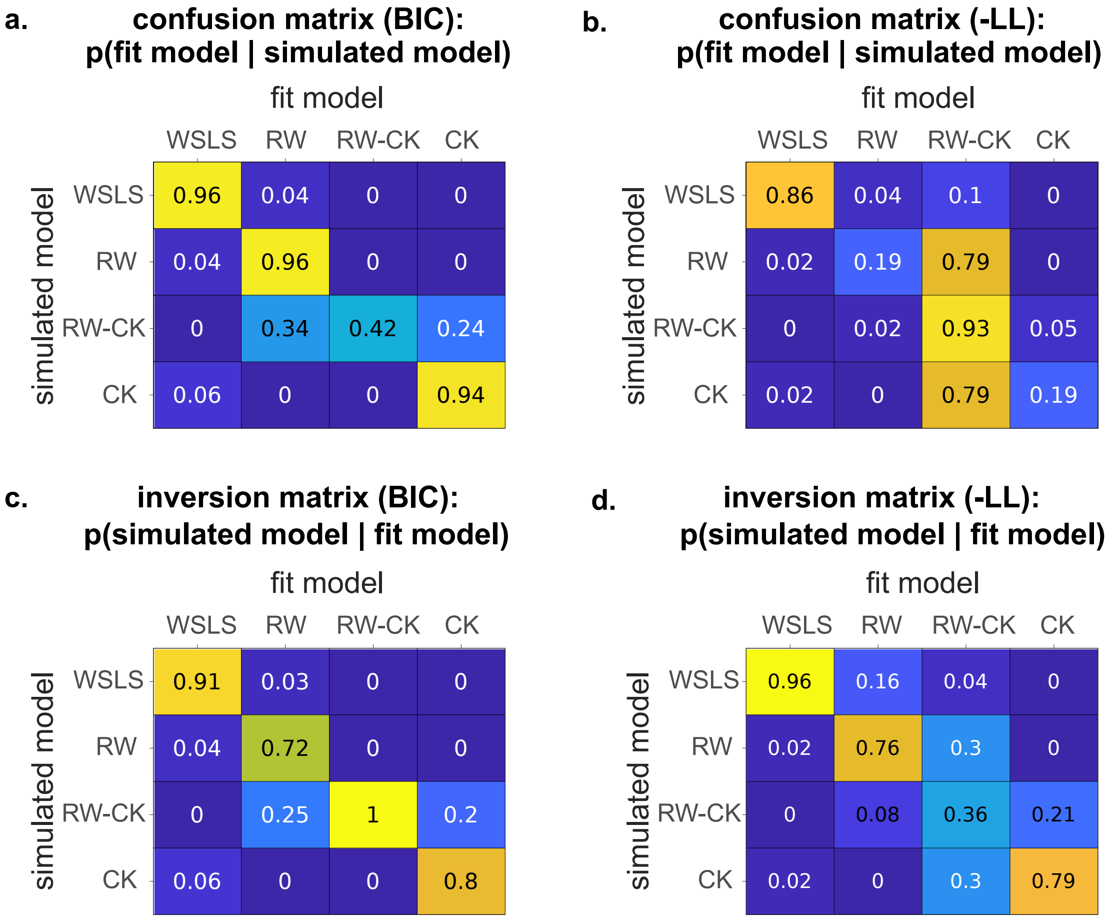


Figure S3: **Model recovery confusion matrices (CM) and inversion matrices (IM). a,** As indicated by a CM derived from BIC values, the data from WSLS, RW, and CK models can be accurately recovered. The combined RW and CK model (RW-CK) shows moderate performance in recovering its ground-truth model, with a portion of the data being better explained by the simple RW (34%) and CK (24%) models. **b,** We tested if RW-CK can best recover its own data when it is not being penalized for its additional free parameters. Minimizing the negative log-likelihood, we find that the RW-CK model recovery improves drastically (93%). As expected, it also captures the data generated by the simple RW (79%) and CK (79%) models. **c,** Additionally, we computed IM based on the CM obtained from BIC (subplot a). The matrix indicates that though the recovery of RW-CK was moderate (42%), we can be certain that it generated the data, given that it fits the data best (100%). **d,** In contrast, our confidence that RW-CK model generated its data (given that it fits the data best) drops to 36% when estimating model recovery using log-likelihood values.

A perfect model recovery would result in an identity matrix. The confusion matrix estimated using the BIC values suggest that the data generated by WSLS, RW, and CK models are best recovered by their own model (Fig. S3a). However, though the RW-CK model moderately recovered its own data, the remaining data generated by this model were being recovered by the simple RW and CK models. This observation may reflect the shared features of the RW-CK model and the individual RW and CK models. Additionally, considering that BIC was employed to compute the confusion matrix, the model recovery of RW-CK might have been impacted by the penalization of the additional two free parameters.

We thus assessed whether the RW-CK model recovery improves by using log-likelihood values instead of BIC values. Indeed, the RW-CK model successfully recovered its own data as well as the data generated by the RW and CK models (Fig. S3b).

As suggested by Wilson & Collins (2019), we also examined the inverse of the confusion matrix, i.e., p(simulated model | fit model). Simply put, the matrix allows us to make the following inference: Given that a model fits data, which model likely generated the data? This inversion matrix offers better interpretability when the true model is unknown, which is often the case when testing the fit of a model to participant data. The matrix is computed from the confusion matrix by Bayes rule and referred to as inversion matrix. The inversion matrix computed with BIC (Fig. S3c) shows a high confidence (100%) that the RW-CK model generated the data given that it fits the data best. By contrast, when using log-likelihood values, the probability is drastically reduced that the RW-CK generated its own data, given that it fits the data best. Consequently, we used the BIC values for analyzing the model fit of the empirical data.

### 3 RW-estimated HR probability correlates with simulation-based liking update

To assess the overall probability of the RW model to select the HR people, we first estimated trial-wise probabilities of selecting a HR person. This was done using the RW-based choice values and the softmax rule (see Eqn. 2). We then computed the area under this curve (AUC) using the approach outlined by Pruessner et al. (2003). Across participants, the AUC correlated with the increase in liking for the HR people as measured on the external rating task (robust skipped Spearman's correlation between AUC and HR relative to LR update: rs = .46, 95% CI = [.20 .66]; robust skipped Spearman's correlation between AUC and HR update: rs = .41, 95% CI = [.16 .60]; Fig. S4).

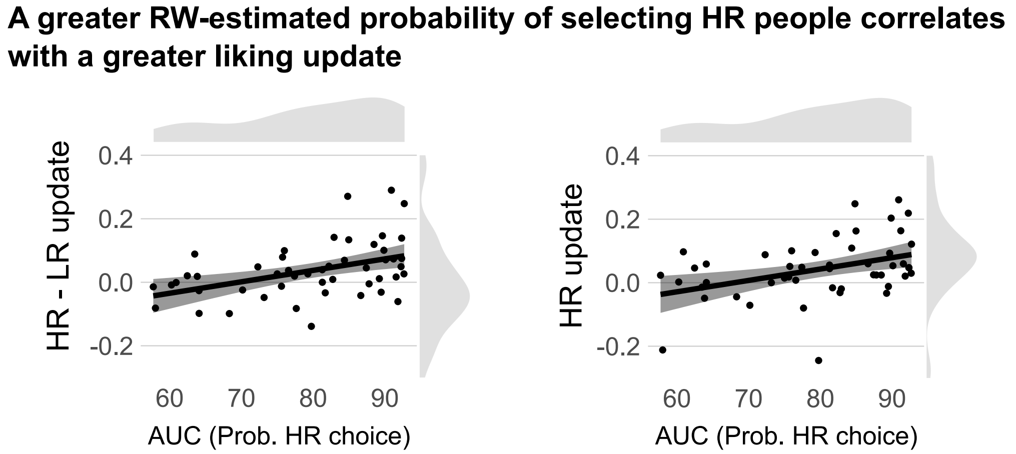


Figure S4: The RW model estimation of HR probability correlated with **a,** the update in liking for HR vs LR people **b,** and also with the update in liking for HR people without adjusting for the update for LR people.

### 4 Prediction-error (PE) parametric-modulation analysis

**Table S1:** Prediction error modulates univariate whole-brain activity.

| AAL Label | Hemisphere | Voxels | x | y | z | t-value |
|---|---|---|---|---|---|---|
| Olfactory[1] | L | 11223 | -4 | 22 | -10 | 12.22 |
| Fusiform[2] | L | | -27 | -33 | -21 | 11.44 |
| Olfactory[1] | R | | 1 | 12 | -7 | 11.07 |
| Precuneus[4] | L | | -7 | -58 | 18 | 10.97 |
| ParaHippocampal[2,3] | L | | -19 | -13 | -21 | 10.94 |
| Fusiform[2,3] | R | | 28 | -38 | -15 | 10.36 |
| Frontal Med Orb[5] | L | | -9 | 47 | -12 | 9.86 |
| Postcentral | L | 8810 | -39 | -30 | 62 | 12.18 |
| Precentral | L | | -27 | -23 | 70 | 10.85 |
| Postcentral | L | | -27 | -40 | 67 | 9.44 |
| Frontal Mid | L | 451 | -24 | 29 | 48 | 8.52 |
| Frontal Sup | L | | -22 | 34 | 34 | 5.67 |
| Cerebelum 8 | L | 517 | -22 | -58 | -56 | 6.67 |
| | | | -19 | -60 | -46 | 6.29 |
| Cerebelum 7b | L | | -12 | -75 | -48 | 6.07 |
| Caudate | L | 46 | -17 | 9 | 23 | 5.88 |
| Frontal Inf Orb | L | 77 | -29 | 34 | -12 | 5.79 |
| Temporal Inf | L | 52 | -54 | -48 | -15 | 5.15 |
| Cerebelum Crus2 | R | 28 | 48 | -68 | -40 | 4.83 |
| Frontal Inf Orb | R | 27 | 28 | 37 | -12 | 4.42 |
| Frontal Sup Orb | R | | 18 | 37 | -18 | 3.99 |
| Frontal Mid | R | 89 | 25 | 29 | 40 | 4.37 |
| | R | | 30 | 39 | 34 | 4.33 |

*Note.* Thresholded at $p < 0.001$, uncorrected; minimum of 20 voxels; [1]cluster contains ventral striatum; [2]cluster contains parahippocampal cortex; [3]cluster contains hippocampus; [4]cluster contains paracingulate cortex including the retrosplenial cortex; [5]cluster contains ventromedial prefrontal cortex.

**Table S2:** Results from small volume correction within the ventral striatum ROI.

| p(FWE-corr) | p(unc) | t-value | x | y | z |
|---|---|---|---|---|---|
| < 0.001 | < 0.001 | 8.49 | 6 | 12 | -7 |
| < 0.001 | < 0.001 | 7.68 | -4 | 7 | -7 |
| < 0.001 | < 0.001 | 7.56 | -7 | 14 | -7 |
| < 0.001 | < 0.001 | 7.30 | -4 | 12 | -4 |
| 0.006 | < 0.001 | 4.33 | -14 | 9 | -12 |

### 5 Choice value (Q) parametric modulation analysis results

**Table S3:** Choice value (Q) modulates univariate whole-brain activity.

| AAL Label | Hemisphere | Voxels | x | y | z | t-value |
|---|---|---|---|---|---|---|
| Cerebelum Crus2 | L | 319 | -24 | -75 | -40 | 7.36 |
| | L | | -34 | -85 | -32 | 5.39 |
| | L | | -22 | -85 | -37 | 5.01 |
| SupraMarginal | R | 2310 | 60 | -35 | 29 | 7.11 |
| | R | | 58 | -26 | 26 | 6.5 |
| Putamen | R | | 30 | -18 | 4 | 6.41 |
| Cerebelum Crus2 | R | 214 | 20 | -75 | -37 | 6.85 |
| | R | | 33 | -85 | -34 | 5.01 |
| Temporal Mid | L | 117 | -54 | 2 | -15 | 5.99 |
| Temporal Pole Sup | L | | -42 | 2 | -15 | 3.59 |
| Cingulum Mid | R | 452 | 8 | -21 | 42 | 5.89 |
| Precuneus | L | | -7 | -53 | 59 | 5.12 |
| Postcentral | L | | -24 | -43 | 67 | 5.05 |
| Insula | L | 844 | -39 | 7 | 1 | 5.93 |
| Putamen | L | | -27 | 4 | 10 | 5.58 |
| | L | | -27 | -8 | 4 | 5.47 |
| Cingulum Mid | R | 466 | 8 | -21 | 42 | 5.92 |
| Precuneus | L | | -7 | -53 | 59 | 5.18 |
| Postcentral | L | | -24 | -43 | 67 | 5.11 |
| SupraMarginal | L | 334 | -67 | -43 | 29 | 5.67 |
| | L | | -62 | -40 | 37 | 5.59 |
| | L | | -67 | -33 | 26 | 5.54 |
| Frontal Mid | L | 147 | -32 | 47 | 26 | 5.35 |
| | L | | -24 | 39 | 26 | 4.58 |
| | L | | -32 | 32 | 45 | 3.91 |
| **Frontal Sup Medial*** | R | 111 | 3 | 62 | 34 | 5.21 |
| | L | | -4 | 52 | 48 | 4.6 |
| Cerebelum 8 | L | 38 | -22 | -65 | -54 | 4.8 |
| Temporal Mid | L | 83 | -62 | -55 | -2 | 4.69 |
| | L | | -57 | -63 | -2 | 4.2 |
| | L | | -54 | -55 | 10 | 3.81 |
| Cingulum Ant | R | 27 | 1 | 24 | 29 | 4.31 |
| Supp Motor Area | R | 93 | 6 | 7 | 70 | 4.18 |
| | R | | 8 | -3 | 64 | 3.85 |
| | R | | 6 | 24 | 64 | 3.79 |

| | | | | | | |
|---|---|---|---|---|---|---|
| Temporal Sup | L | 26 | -57 | -28 | 10 | 4.18 |
| Frontal Mid | R | 22 | 28 | 54 | 23 | 4.02 |
| Frontal Sup | L | 27 | -19 | -1 | 70 | 3.66 |
| Supp Motor Area | L | | -9 | -8 | 70 | 3.59 |

*Note.* Thresholded at $p < 0.001$, uncorrected; minimum of 20 voxels; *cluster contains dmPFC ROI.

**Table S4:** Results from small volume correction within the dmPFC ROI.

| p(FWE-corr) | p(unc) | t-value | x | y | z |
|---|---|---|---|---|---|
| 0.002 | < 0.001 | 4.33 | 10 | 62 | 29 |

#### 6 Searchlight RSA results

We used searchlight RSA analysis to identify brain regions that encode representations of individual people (sphere radius of 7.5 mm, 3 voxels). We used functions from the RSA toolbox (Nili et al., 2014) and compared activity patterns across runs in a similar fashion as done for the ROI-based RSA. Before running the second-level analysis, first-level maps were z-transformed, normalized to MNI space using the fMRIprep transform files, and smoothed with a Gaussian Kernel of 6 mm radius of FWHM.

**Table S5:** Results of the searchlight representational-similarity analysis.

| AAL Label | Hemisphere | Voxels | x | y | z | t-value |
|---|---|---|---|---|---|---|
| Occipital Sup | L | 10119 | -22 | -73 | 37 | 4.47 |
| | L | | -26 | -68 | 31 | 4.25 |
| | L | | -29 | -86 | 30 | 4.13 |
| Temporal Pole Mid | R | 1660 | 41 | 11 | -34 | 4.46 |
| Temporal Mid | L | 4706 | -42 | -54 | 14 | 4.29 |
| | L | | -51 | -56 | 9 | 4.2 |
| | L | | -47 | -49 | 2 | 3.72 |
| Cingulum Mid | L | 2245 | -5 | -25 | 39 | 4.21 |
| | R | | 4 | -35 | 37 | 3.8 |
| | L | | -9 | -22 | 47 | 3.67 |
| Occipital Mid | R | 2140 | 38 | -78 | 26 | 4.18 |
| Frontal Sup | L | 1346 | -19 | 38 | 40 | 4.1 |
| Frontal Mid | L | | -28 | 31 | 49 | 3.48 |
| Precuneus | R | 1917 | 16 | -47 | 13 | 4.08 |
| Cingulum Post | R | | 6 | -39 | 7 | 3.69 |
| Precuneus | R | 1098 | 16 | -58 | 35 | 4.08 |
| | R | | 27 | -56 | 28 | 3.46 |

| | | | | | | |
|---|---|---|---|---|---|---|
| Fusiform | R | 466 | 39 | -37 | -13 | 4.05 |
| Fusiform | L | 1757 | -25 | -41 | -17 | 4.02 |
| | L | | -34 | -30 | -21 | 3.38 |
| Cingulum Post | L | 2071 | -9 | -45 | 24 | 4.01 |
| Precuneus | L | | 0 | -57 | 32 | 3.95 |
| Putamen | R | 300 | 28 | 0 | 9 | 3.97 |
| Caudate | L | 273 | -27 | -11 | 32 | 3.92 |
| Putamen | L | 313 | -29 | 3 | 1 | 3.91 |
| Cingulum Ant | L | 830 | -16 | 43 | 12 | 3.87 |
| | L | | -8 | 41 | 12 | 3.57 |
| Cerebelum 10 | L | 214 | -11 | -29 | -39 | 3.79 |
| **Frontal Sup Medial*** | R | 643 | 6 | 60 | 21 | 3.75 |
| Precentral | L | 83 | -29 | -10 | 75 | 3.68 |
| Frontal Sup Orb | R | 236 | 21 | 23 | -11 | 3.64 |
| Thalamus | L | 79 | -19 | -19 | 4 | 3.63 |
| Hippocampus | L | 145 | -13 | -36 | 4 | 3.61 |
| Hippocampus | R | 205 | 39 | -17 | -10 | 3.57 |
| Frontal Med Orb | L | 129 | -1 | 52 | -8 | 3.53 |
| Frontal Mid | L | 290 | -28 | 49 | 22 | 3.53 |
| Precuneus | L | 160 | -2 | -52 | 57 | 3.52 |
| Frontal Sup | L | 292 | -18 | 23 | 60 | 3.5 |
| Frontal Inf Orb | R | 47 | 42 | 39 | -8 | 3.47 |
| Putamen | R | 42 | 30 | 15 | -2 | 3.42 |
| Frontal Sup Medial | L | 176 | -9 | 51 | 32 | 3.4 |
| Olfactory | R | 22 | 6 | 20 | -7 | 3.38 |

*Note.* Thresholded at $p < 0.001$, uncorrected; minimum of 20 voxels; *cluster contains dmPFC ROI.

**Table S6:** Results from small volume correction within the dmPFC ROI.

| p(FWE-corr) | p(unc) | t-value | x | y | z |
|---|---|---|---|---|---|
| 0.007 | < 0.001 | 3.73 | 6 | 60 | 20 |
| 0.019 | 0.001 | 3.33 | 7 | 65 | 21 |

### 7 Psychophysiological interaction (PPI) analysis

**Table S7:** Brain regions exhibiting PE-dependent coupling with the ventral striatum.

| AAL Label | Hemisphere | Voxels | x | y | z | t-value |
|---|---|---|---|---|---|---|
| **Frontal Sup Medial*** | L | 650 | -7 | 62 | 32 | 7.91 |
| | R | | 3 | 62 | 29 | 5.92 |
| | L | | -9 | 59 | 15 | 5.23 |
| Temporal Pole Sup | L | 1877 | -39 | 22 | -29 | 7.2 |
| Frontal Inf Orb | L | | -47 | 32 | -2 | 7.03 |
| Frontal Inf Tri | L | | -59 | 24 | 10 | 6.61 |
| Temporal Pole Sup | R | 423 | 48 | 22 | -26 | 7.19 |
| Temporal Sup | R | | 58 | 4 | -12 | 5.87 |
| Frontal Inf Orb | R | | 38 | 39 | -7 | 5.68 |
| Supp Motor Area | L | 736 | -4 | 12 | 54 | 7.06 |
| | L | | -2 | 2 | 64 | 6.09 |
| | R | | 6 | 12 | 64 | 5.53 |
| Frontal Mid | L | 314 | -42 | 4 | 51 | 6.51 |
| Precentral | L | | -52 | -1 | 48 | 4.74 |
| Frontal Sup | L | | -24 | -3 | 56 | 4.41 |
| Occipital Inf | L | 732 | -44 | -73 | -15 | 6.5 |
| Calcarine | L | | -12 | -103 | -10 | 5.99 |
| Lingual | L | | -32 | -95 | -15 | 5.91 |
| Calcarine | R | 502 | 18 | -100 | -4 | 6.34 |
| Occipital Inf | R | | 33 | -95 | -4 | 5.58 |
| Lingual | R | | 23 | -90 | -12 | 5.45 |
| Precuneus | L | 371 | -4 | -55 | 10 | 6.2 |
| Calcarine | L | | -14 | -48 | 7 | 5.91 |
| Precuneus | L | | -9 | -58 | 20 | 5.09 |
| Frontal Inf Orb | R | 118 | 53 | 32 | -2 | 6.07 |
| Olfactory | R | 445 | 1 | 17 | -7 | 6.06 |
| Rectus | L | | -2 | 49 | -21 | 5.8 |
| Frontal Med Orb | L | | -2 | 64 | -10 | 5.78 |
| Hippocampus | R | 100 | 25 | -13 | -12 | 5.73 |
| Fusiform | R | | 25 | -28 | -18 | 4.65 |
| Hippocampus | R | | 18 | -21 | -10 | 3.78 |
| Angular | L | 133 | -52 | -75 | 26 | 5.68 |
| Occipital Mid | L | | -39 | -80 | 37 | 5.4 |
| | L | | -37 | -65 | 26 | 3.45 |
| Hippocampus | L | 128 | -24 | -16 | -12 | 5.3 |

| | | | | | | |
|---|---|---|---|---|---|---|
| | L | | -34 | -11 | -15 | 5.05 |
| | L | | -19 | -6 | -15 | 4.67 |
| Cerebelum 9 | R | 64 | 8 | -48 | -46 | 5.2 |
| Temporal Mid | L | 27 | -57 | -48 | 23 | 4.92 |
| Caudate | L | 44 | -14 | 12 | 12 | 4.6 |
| Pallidum | L | | -14 | 4 | -4 | 3.6 |
| Putamen | L | | -19 | 9 | 1 | 3.51 |
| Precentral | R | 51 | 38 | -23 | 51 | 4.47 |
| | R | | 30 | -28 | 59 | 3.79 |
| Cerebelum Crus2 | R | 46 | 15 | -83 | -43 | 4.44 |
| Cerebelum 8 | R | 36 | 35 | -60 | -54 | 4.33 |
| | R | | 25 | -68 | -51 | 4.02 |
| Cerebelum Crus1 | R | 45 | 38 | -60 | -34 | 4.11 |
| | R | | 35 | -63 | -26 | 3.97 |
| Temporal Sup | R | 23 | 48 | -23 | 1 | 3.89 |
| | R | | 43 | -33 | 4 | 3.73 |
| | R | | 53 | -30 | 4 | 3.62 |

*Note.* Thresholded at $p < 0.001$, uncorrected; minimum of 20 voxels; *cluster contains dmPFC ROI.

**Table S8:** Results from small volume correction within the dmPFC ROI.

| p(FWE-corr) | p(unc) | t-value | x | y | z |
|---|---|---|---|---|---|
| < 0.001 | < 0.001 | 5.61 | 3 | 62 | 26 |
| < 0.001 | < 0.001 | 5.24 | 3 | 67 | 26 |
| 0.030 | 0.001 | 3.4 | 6 | 57 | 15 |

### 8 References

Nili, H., Wingfield, C., Walther, A., Su, L., Marslen-Wilson, W., & Kriegeskorte, N. (2014). A Toolbox for Representational Similarity Analysis. *PLOS Computational Biology*, 10(4), e1003553. https://doi.org/10.1371/journal.pcbi.1003553

Pruessner, J. C., Kirschbaum, C., Meinlschmid, G., & Hellhammer, D. H. (2003). Two formulas for computation of the area under the curve represent measures of total hormone concentration versus time-dependent change. *Psychoneuroendocrinology*, 28(7), 916–931. https://doi.org/10.1016/S0306-4530(02)00108-7

Wilson, R. C., & Collins, A. G. (2019). Ten simple rules for the computational modeling of behavioral data. *eLife*, 8, e49547. https://doi.org/10.7554/eLife.49547
